# Physiological variability in mitochondrial rRNA predisposes to metabolic syndrome

**DOI:** 10.1101/2023.11.08.566188

**Authors:** Petr Pecina, Kristýna Čunátová, Vilma Kaplanová, Guillermo Puertas-Frias, Jan Šilhavý, Kateřina Tauchmannová, Marek Vrbacký, Tomáš Čajka, Ondřej Gahura, Michal Pravenec, Josef Houštěk, Tomáš Mráček, Alena Pecinová

## Abstract

Metabolic syndrome is a growing concern in developed societies, and due to its polygenic nature, the genetic component is only slowly being elucidated. Common mitochondrial DNA sequence variants have been associated with symptoms of metabolic syndrome and may be relevant players in the genetics of metabolic syndrome.

We investigate the effect of mitochondrial sequence variation on the metabolic phenotype in conplastic rat strains with identical nuclear but unique mitochondrial genomes, challenged by high-fat diet. We find that the variation in mitochondrial rRNA sequence represents a risk factor in insulin resistance development, which is caused by diacylglycerols accumulation induced by tissue-specific reduction of the oxidative capacity. These metabolic perturbations stem from the 12S rRNA sequence variation affecting mitochondrial ribosome assembly and translation. Our work demonstrates that physiological variation in mitochondrial rRNA might represent a relevant underlying factor in the progression of metabolic syndrome.

**Competing interests:** The authors declare that no competing interests exist.

## Introduction

Mitochondria, the organelles of bacterial origin, still maintain mitochondrial DNA (mtDNA), and its sequence exerts variability (single nucleotide polymorphisms - SNPs), dividing the population into mtDNA families known as haplogroups (Kenney et al. 2014). Unlike nuclear DNA (nDNA), mtDNA in mammals is maternally inherited and encodes only 13 structural proteins, the subunits of the oxidative phosphorylation apparatus (OXPHOS), two ribosomal RNAs and 22 transfer RNAs required for mitochondrial protein synthesis. The nuclear genome encodes the remaining circa 1500 mitochondrial proteins, which are transported into the mitochondria (Shadel and Clayton 1997). This requires coordination in expression and translation between the two genomes (Cagin and Enriquez 2015, Dennerlein et al. 2017). Recently, studies in human populations have associated mtDNA haplogroups with the risk of metabolic syndrome or its complications (Chinnery et al. 2010, Estopinal et al. 2014, Martikainen et al. 2015), and sequence variation in mtDNA has also been shown to influence the expression of stress response genes (Bellizzi et al. 2006).

A higher risk of metabolic syndrome is strongly associated with weight gain due to high nutrient excess (Grundy 2004). The resulting aberrant distribution of fat, mainly in visceral and intra-abdominal depots (Vazquez et al. 2007, Castro et al. 2014), leads to lipotoxicity – ectopic fat storage in metabolically active tissues such as the liver, myocardium, or skeletal muscle (Unger 2002). The metabolic disbalance is associated with symptoms such as insulin resistance, type 2 diabetes mellitus (T2DM), non-alcoholic fatty liver disease, or cardiovascular disease (reviewed in (Boutari and Mantzoros 2022)).

While the link between mitochondrial function and metabolic syndrome is widely accepted, the molecular mechanisms behind it still need to be better defined. Many studies have implicated a decrease in mitochondrial oxidative capacity as a primary cause of insulin resistance and T2DM (Kelley et al. 1999, Mootha et al. 2003)), resulting in the accumulation of bioactive lipids and increased risk of T2DM. Inefficient fatty acid oxidation and subsequent accumulation of bioactive lipids such as diacylglycerols (DGs), fatty acyl-CoA, or ceramides have been shown to inhibit the insulin signalling pathway (Yu et al. 2002, Chavez and Summers 2012). Specifically, it has been observed that an increase in plasma fatty acids results in elevated intracellular fatty acyl-CoA and DGs in skeletal muscle, leading to activation of protein kinase C θ (PKCθ). Subsequent phosphorylation of IRS-1 prevents insulin-stimulated tyrosine phosphorylation and decreases insulin-stimulated glucose transport (Yu et al. 2002). In contrast to DGs accumulation, high ceramide concentrations directly affect the activity of the serine/threonine protein kinase Akt/PKB, either through activation of phosphatase 2A (Teruel et al. 2001, Chavez and Summers 2012) or through protein kinase C ζ (PKCζ) (Powell et al. 2003).

Another possible mechanism linking mitochondrial dysfunction to insulin resistance is the mitochondrial generation of reactive oxygen species (ROS). An increased supply of electrons from excess nutrients increases the likelihood of their slip to molecular oxygen, forming superoxide and, subsequently, other forms of ROS, which may directly induce insulin resistance, or damage mitochondrial proteins, lipids or DNA, thereby reducing oxidative capacity (Anderson et al. 2009).

Insulin resistance has also been associated with low-grade inflammation in obese and diabetic patients (Pickup and Crook 1998, Wu and Ballantyne 2020, Jani et al. 2021). This condition is characterised by chronically elevated inflammatory markers such as TNF-α, interleukin-1β or interleukin-6 (Tornatore et al. 2012). Also, mice overexpressing the chemokine ligand CCL2 (also known as MCP1) in adipose tissue exhibit insulin resistance (Kanda et al. 2006). Moreover, elevated cytokine levels correlated with increased lipolysis in adipose tissue, suggesting destabilised lipid droplets as a possible mechanism allowing better access of lipases to triacylglycerols (reviewed in (Samuel and Shulman 2012)).

In the current project, we asked whether the physiological variation in the mitochondrial DNA sequence may directly contribute to symptoms of metabolic syndrome. For this purpose, we used our unique model of conplastic rats carrying mtDNA from spontaneously hypertensive rat strain (mtSHR), Brown Norway strain (mtBN) or Fischer strain (mtF344) on the identical nuclear background (SHR) (Pravenec et al. 2007, Houstek et al. 2012, Houstek et al. 2014, Pravenec et al. 2021). We found that mtF344 animals develop insulin resistance on a high-fat diet, which can be explained by differences in mitochondrial 12S rRNA leading to reduced substrate oxidation and subsequent accumulation of bioactive lipids and insulin resistance.

## Results

Our previous work has established and characterised rat conplastic strains, which differ only in their mtDNA (Pravenec et al. 2007, Houstek et al. 2012, Houstek et al. 2014). Numerous SNP variants exist in mtDNA across the strains (summarised in Supplementary tables S1, S2). Overall, they affect 8 genes encoding for subunits of oxidative phosphorylation complexes, 7 tRNAs, and 12S as well as 16S rRNAs. As our previous data indicated that conplastic strains differ in their metabolic phenotypes, we asked to what extent mtDNA sequence variability may affect insulin sensitivity and whether a high-fat diet could further aggravate such phenotype. Therefore, we designed an experiment to compare SHR rats (mtSHR) with conplastic strains possessing mtDNA from F344 or BN strain on the SHR nuclear background (mtF344 or mtBN, respectively). After weaning (week 5), the animals were transferred to either chow (CHD) or high-fat (HFD) diet for 15 weeks. The oral glucose tolerance test (OGTT) was performed at week 18 (2 weeks before experiment termination and tissue collection) to assess insulin sensitivity. The whole experimental design is summarised in Figure 1A.

**Figure 1.**
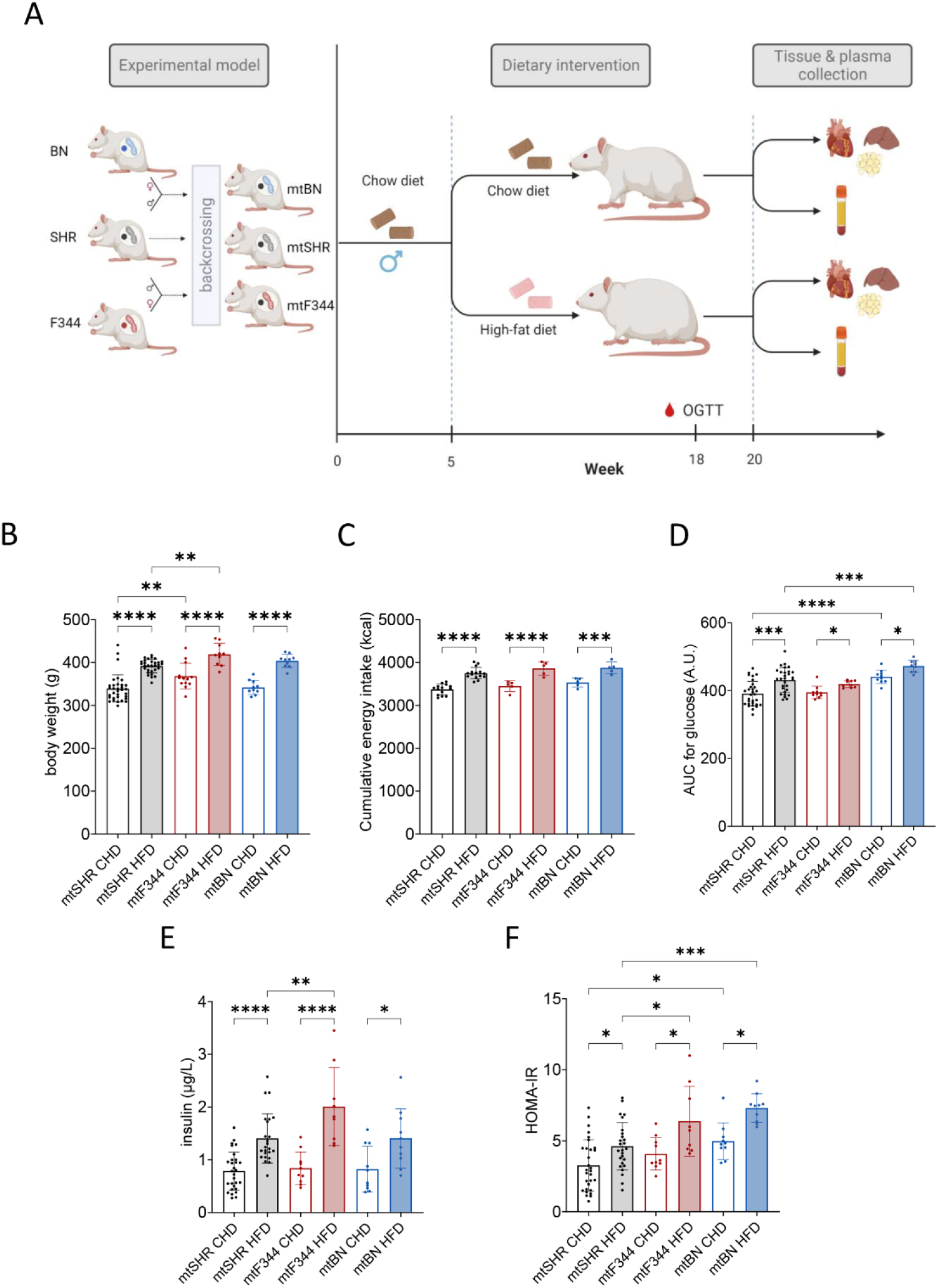
Experimental design and metabolic phenotype of conplastic rats. (A) Schematic depicting of conplastic strains created by multiple backcrossing of males SHR rats with females of SHR, and females of progeny derived from (♀BN x ♂SHR)F1 or (♀F344 x ♂SHR)F1 hybrids. After weaning, experimental groups were transferred to a chow or high-fat diet for 15 weeks. At week 18, the OGTT test was performed, and at week 20, the tissues were collected for further analyses. Body weights (B) and cumulative energy intake (C) after 15 weeks of dietary intervention. (D) Area under the curve calculated from oral glucose tolerance test. (E) Insulin levels 30 min after glucose gavage and (F) homeostatic model assessment (HOMA-IR) calculated from fasting glucose and insulin levels. Data represent means ± S.D. from at least 9 animals. Asterisks represent p-value: * <0.05; ** <0.01; *** <0.001 **** <0.0001.

### The high-fat diet promotes insulin resistance in conplastic rats

As presumed, the HFD diet led to an increase in mean body weight in all the strains. Relative to mtSHR strain, the body weight of mtF344 was increased on either of the diets (Figure 1B), and this was not caused by increased calorie intake (Figure 1C). As expected, the area under the curve (AUC) for glucose during OGTT was higher on HFD compared to CHD in all groups, indicating a lower ability to clear glucose from the blood (Figure 1D). Also, insulin levels were increased significantly 30 min after glucose gavage on HFD (Figure 1E). Interestingly, while AUC for glucose did not differ in mtF344 strain compared to mtSHR on either of the diets, insulin concentration was significantly higher in mtF344 animals on HFD. On the other hand, in the mtBN strain, glucose tolerance was decreased, while insulin levels remained unchanged (Figure 1D, E).

We calculated homeostatic model assessment (HOMA-IR) to quantify insulin sensitivity from fasting glucose and insulin levels (Figure 1F). As expected, HFD led to a significant increase in HOMA-IR in all groups compared to CHD. Importantly, we also observed a significant genotype-dependent increase of HOMA-IR in mtF344 animals on HFD and in mtBN on both diets. These data indicate that relative to mtSHR, both mtF344 and mtBN mtDNA variants predispose to impaired insulin sensitivity on a high-fat diet.

### The high-fat diet-induced insulin resistance in mtF344 correlates with the accumulation of diacylglycerols

As a next step, we focused on the possible mechanism(s) responsible for differences in insulin sensitivity between conplastic animals. In principle, three different mechanisms were proposed to link mitochondrial function and obesity-induced insulin resistance: i) elevated levels of inflammatory cytokines, ii) increased oxidative stress, iii) accumulation of bioactive lipids such as ceramides or DGs resulting from reduced oxidative capacity of mitochondria.

First, we focused on inflammatory response and assessed pro-inflammatory cytokine Il1b and chemokine Ccl2 levels in the liver and white adipose tissue. As expected, the expression of the genes was increased in animals on HFD (Figures 2A, B, D and E), and the concentration of Il1b, but not Ccl2, measured by multiplex immunoassay was also increased on HFD in the liver (Figures 2C, F). However, no difference between genotypes was observed in expression profiles. In summary, the changes in inflammatory response do not explain the strain-specific differences in insulin sensitivity.

**Figure 2.**
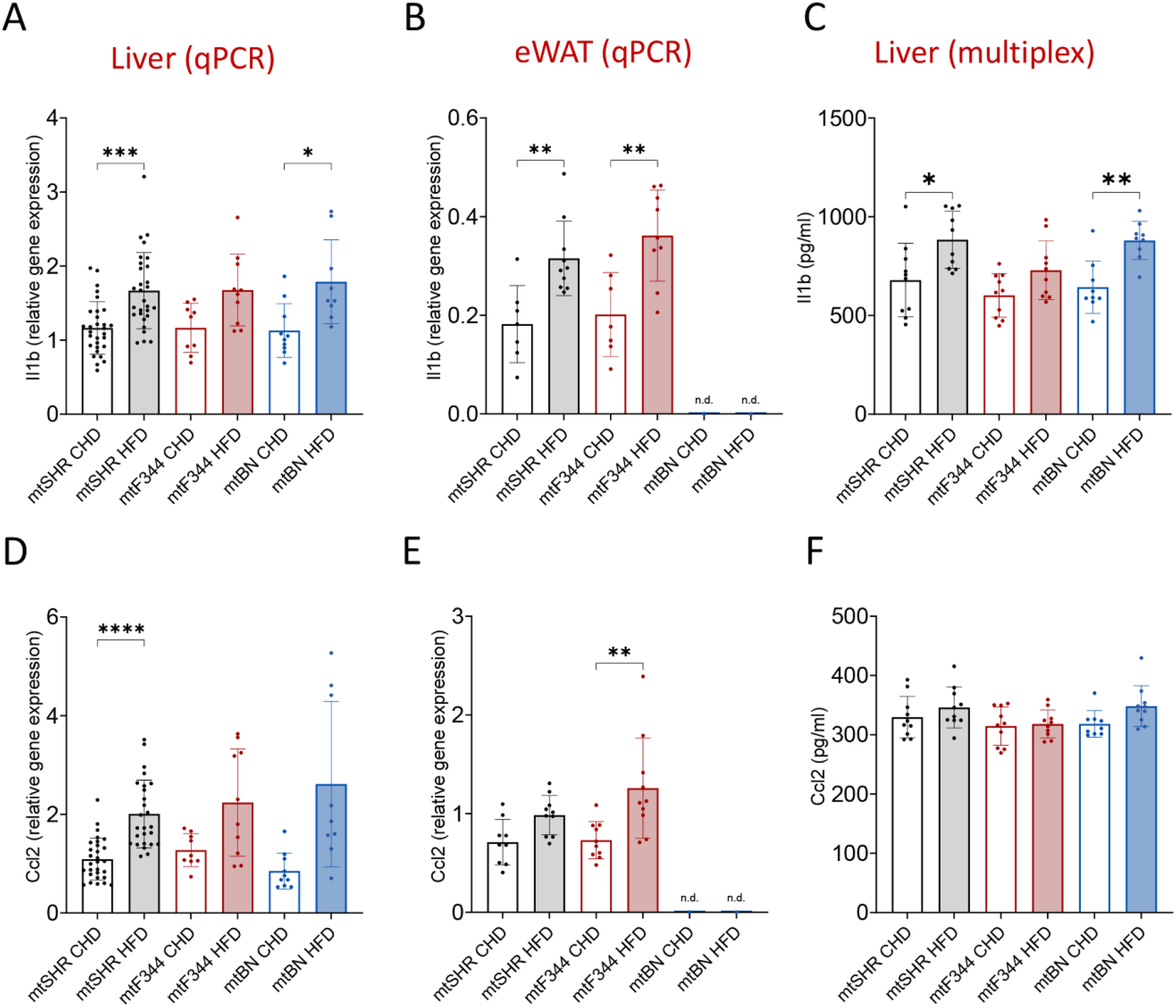
Inflammatory response in conplastic rats. Gene expression of pro-inflammatory cytokine *Il1b* (A, B) and chemokine *Ccl2* (D, E) in the liver (A, D) or epididymal adipose tissue (eWAT, B, E). Corresponding Il1b (C) and Ccl2 (F) levels in liver extracts measured by bead-based immunoassay. Data represent means ± S.D. from at least 7 animals. Asterisks represent p-value: * <0.05; ** <0.01; ***<0.001 **** <0.0001.

Since both nuclear and mitochondrial genomes encode the subunits of mitochondrial oxidative phosphorylation complexes, their sterical inconsistency may affect the stability of the complexes and, consequently, increase the production of reactive oxygen species (ROS). Therefore, we measured H_2_O_2_ production by Amplex red assay in isolated rat liver mitochondria. To distinguish where the electrons may escape towards oxygen, we used a combination of specific substrates and inhibitors of respiratory chain dehydrogenases (Figure 3A). Using NADH-linked substrate (glutamate and malate) and rotenone, we determined the ROS generation on the flavin site of complex I (I_F_, Figure 3B). Compared to mtSHR, ROS production was unchanged in mtBN strain and was slightly lower in mtF344 on CHD.

**Figure 3.**
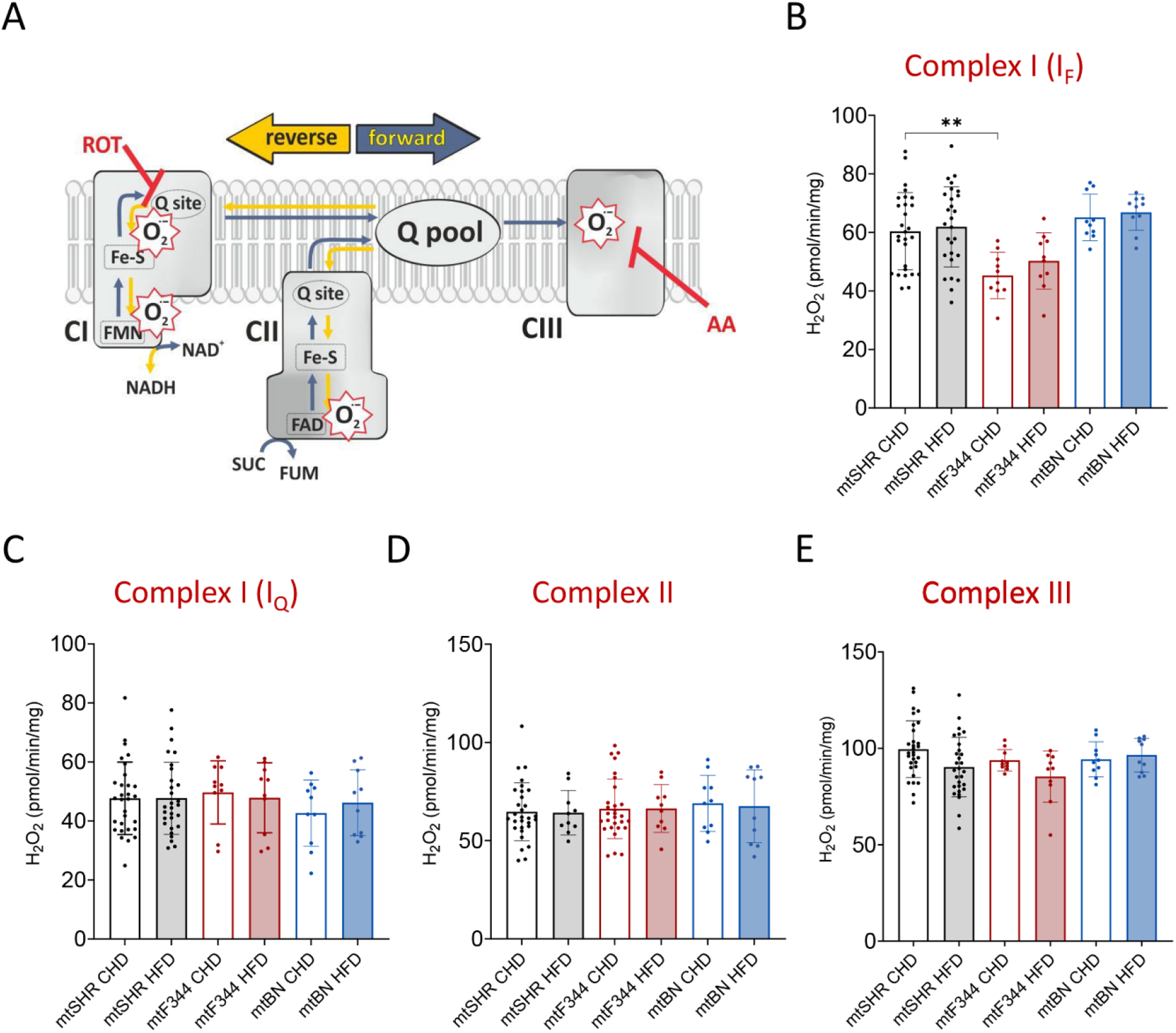
ROS generation in conplastic rats. (A) Scheme of superoxide production in electron transport chain (CI – complex I; CII – complex II; CIII – complex III; ROT – rotenone, complex I inhibitor; AA – antimycin A, complex III inhibitor; O_2_^.-^ - superoxide). (B-E) Production of H_2_O_2_ by isolated liver mitochondria using Amplex Red assay, and specific substrates and inhibitors to distinguish ROS production site. Isolated mitochondria were supplemented with 10 mM glutamate, 2 mM malate and 1 μM rotenone (B), 10 mM succinate and 1 μM rotenone (C), 0.4 mM succinate (D) or 10 mM succinate plus 1 μg/mL antimycin A (E). Data represent means ± S.D. from at least 9 animals. Asterisks represent p-value: * <0.05; ** <0.01; *** <0.001 **** <0.0001.

Under the conditions of high flux from succinate oxidation and high proton-motive force, the electron backflow to complex I occurs. Under such conditions, electrons can escape to molecular oxygen at the Q site (I_Q_), which may be prevented by rotenone (Korshunov et al. 1997, Lambert and Brand 2004). Using saturating succinate concentration (10 mM) and complex I inhibitor rotenone (1 μM), we assessed rotenone-sensitive ROS production. We did not observe any changes in ROS generation between strains or diets (Figure 3C).

Further, we measured ROS production on the flavin of complex II (Figure 3D) using a physiological concentration (0.4 mM) of succinate. As expected, since complex II subunits are encoded solely by nDNA, we did not detect any changes in ROS production between strains. Similarly, no differences were detected with succinate and an inhibitor of complex III (antimycin A), which would reveal the changes in ROS production on complex III (Figure 3E). These data indicate that mitochondria-derived oxidative stress cannot explain the strain-specific differences in insulin sensitivity.

Ultimately, we focused on the content of bioactive lipids, such as ceramides or diacylglycerols, which have also been shown to modulate insulin signalling. For this purpose, we performed unbiased lipidomic profiling of plasma, liver, and heart in all strains on HFD. Lipidomic profiles showed distinguished clusters on a sparse Partial Least Squares Discriminant Analysis (sPLSDA) for individual conplastic strains (Figure 4A). We did not find any effect of haplotype on the levels of ceramides. Relative to mtSHR, the ceramides were even decreased - in all analysed matrices for the mtF344 and in heart only for mtBN (Figure 4B, C). However, in the mtBN strain, we observed a slight but significant increase of triacylglycerols (TGs) in the liver and heart and a slight increase of diacylglycerols (DGs, Figure 4C). Interestingly, in mtF344 animals, DGs were elevated not only in the heart but also in the liver and plasma (Figure 4B). The most abundant DGs, especially in plasma, contain acyl chains that are the most abundant in the diet (Figure S1) – oleic acid (18:1), palmitic acid (16:0), and linoleic acid (18:2). Thus, it is likely that fatty acids stored in DGs originate from diet and did not undergo subsequent remodelling. The results suggest that DGs may serve as modulators of insulin signalling in mtF344 animals.

**Figure 4.**
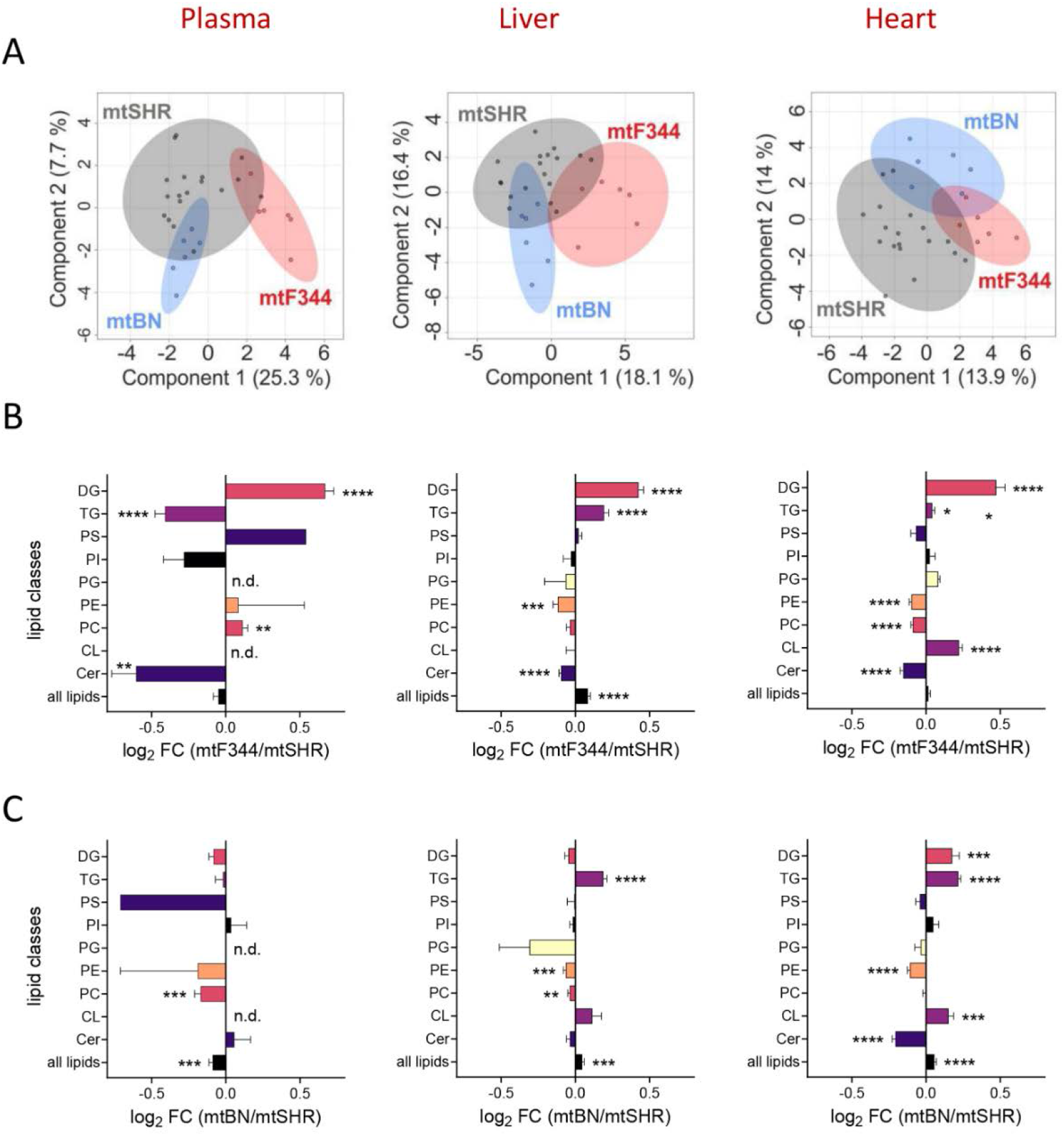
Analysis of lipid classes in conplastic strains on HFD. (A) sPLSDA was performed to differentiate between mtF344, mtBN and mtSHR strains. Each data point represents one animal. (B, C) LC-MS analysis of different lipid classes in plasma, liver, and heart of mtF344 (B) and mtBN (C) strain compared to mtSHR control group. The data are expressed as mean ± SEM of log_2_ fold change from 6 animals. (Lipid classes annotation: Cer, ceramides; CL: cardiolipins; PC: phosphatidylcholines; PE: phosphatidylethanolamines; PG: phosphatidylglycerols; PI: phosphatidylinositols; PS: phosphatidylserines; TG: triacylglycerols; DG: diacylglycerols). The significances were calculated as one sample t-test compared to the mtSHR control group (n = 6), and asterisks represent p-value: * <0.05; ** <0.01; *** <0.001 **** <0.0001.

### mtF344 rats on HFD have lower mitochondrial oxidative capacity in the heart

We hypothesised that DGs accumulation may arise from defective mitochondrial fatty acid oxidation. Therefore, we focused on mitochondrial function and measured mitochondrial respiration in tissue homogenates (Figure 5A). The capacity to oxidise fatty acids was assessed with palmitoyl-carnitine and malate as substrates – a significant HFD-induced increase in fatty acid oxidation capacity (FAO capacity) was observed in the liver of all strains (Figure 5B). Total phosphorylating capacity (OXPHOS capacity) was then measured with a combination of complex I and II substrates, and also tended to be higher after HFD but became statistically significant only in the mtSHR group (Figure 5C). Ultimately, the maximal electron transport chain (ETC) capacity was comparable between genotypes and diets (Figure 5D). The data suggest that liver mitochondria possess a relatively high spare capacity of ETC independent of the mtDNA genotype and may utilise this capacity when fatty acid abundance increases. Therefore, a specific increase in fatty acid oxidation capacity should arise from the upregulation of enzymes involved in mitochondrial β oxidation. Indeed, LFQ proteomic analysis of liver tissue confirmed that many proteins involved in fatty acid oxidation are increased, especially the enzymes required to oxidise long chain fatty acids while there is no profound change in the content of OXPHOS protein subunits (Figure S2).

**Figure 5.**
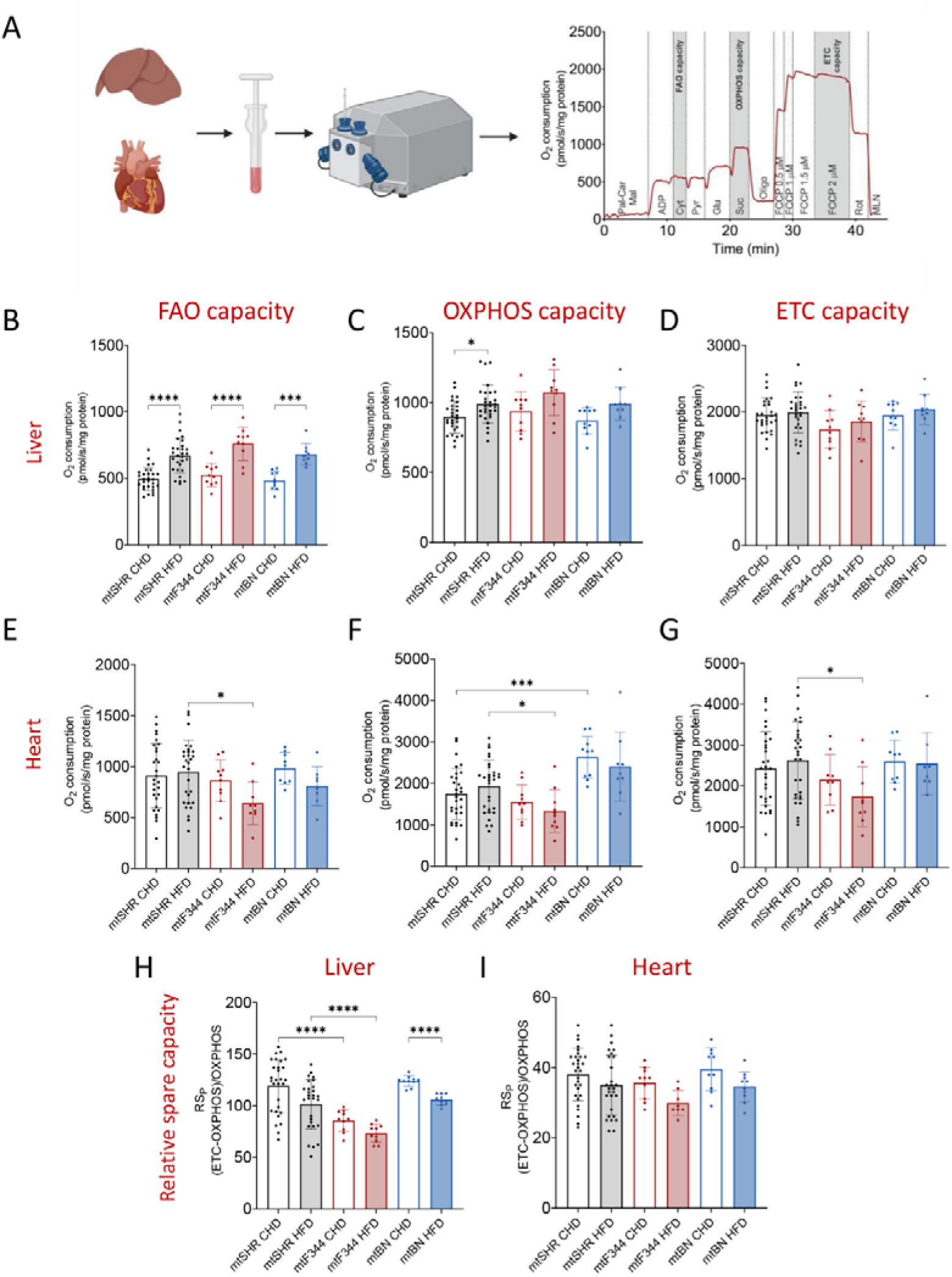
Mitochondrial respiration in conplastic strains. (A) Experimental workflow of the mitochondrial oxygen consumption in the liver and heart homogenates measured by Oxygraph-2k (Oroboros). The representative curve depicts the subsequent addition of different substrates and inhibitors (for more details, see Methods). The liver (B, C) and the heart (E, F) fatty acid oxidation (FAO) and OXPHOS capacities were measured in the presence of 1 mM ADP and either 50 μM palmitoyl carnitine and 2 mM malate (FAO capacity, B and E) or in the combination with 10 mM pyruvate and glutamate (OXPHOS capacity, C and F). ETC capacity (D, G) was determined by combining all substrates in the presence of uncoupler FCCP (1 μM). Relative spare capacity of ETC to OXPHOS capacity (RS_P_) in the liver (H) and heart (I) was calculated as percentage of OXPHOS capacity, i.e. 100*(ETC – OXPHOS capacity)/ OXPHOS capacity. The data are expressed as means ± S.D. from at least 8 animals. Asterisks represent p-value: * <0.05; ** <0.01; *** <0.001 **** <0.0001.

Such diet-dependent phenotype was not observed in the heart, where FAO capacity and overall OXPHOS capacity did not differ between diets in any of the groups (Figure 5E-G). Accordingly, neither the levels of proteins involved in fatty acid metabolism were changed by HFD in mtSHR or mtF344 (Figure S3). In contrast, we observed genotype-specific differences in oxidative capacity. Specifically, in mtBN on CHD, OXPHOS capacity was significantly increased. More strikingly, there was a consistent drop in all evaluated parameters (i.e. FAO and overall OXPHOS capacity as well as total ETC capacity) in mtF344 animals on HFD (Figure 5E-G).

These data imply that HFD represents a stress factor for the mtF344 strain, leading to decreased substrate oxidation in both coupled (FAO/OXPHOS capacity) and uncoupled (ETC capacity) states. Also, in comparison to liver mitochondria, which are metabolically flexible, the heart mitochondria cannot further increase fatty acid oxidation capacity. This implies that mtF344 strain utilises a significant portion of ETC capacity for ADP phosphorylation already on CHD. To compare reserve capacity of the respiratory chain (available for the higher flux demand) between tissues, we assessed the relative spare capacity (RS_p_), which represents a ratio between spare capacity (i.e., ETC – OXPHOS) relative to OXPHOS capacity. RS_P_ capacity in the liver was even higher than OXPHOS capacity (approximately 120 % of OXPHOS capacity). It decreased after HFD (significantly only in the mtBN), which is in agreement with increased flux from fatty acid oxidation and the subsequent higher requirement to utilise the ETC capacity (Figure 5H). RS_P_ capacity was lower in mtF344 strain on both diets (approximately 80% of OXPHOS capacity), yet still sufficient to yield an increase in oxidative capacity on HFD.

On the other hand, the RS_P_ in the heart was considerably lower than in the liver (approximately 40% of OXPHOS capacity) and was practically independent of diet (Figure 5I). This suggests that in the heart response to changes in substrate availability cannot stem from adjustments in the respiratory chain. The unchanged RS_P_ in the mtF344 group reflects that HFD led to a decrease in both OXPHOS and ETC capacities.

### Glucose intolerance in mtBN strain on HFD associates with a selective decrease of complex IV

Although the exposition to HFD led to glucose intolerance in the mtBN strain, it did not alter mitochondrial function in the heart or liver. Analogous phenotype had already been described previously for the mtBN animals on a high-fructose diet (Pravenec et al. 2007). Under these conditions, the phenotype had been ascribed to selective reduction in complex IV (cytochrome *c* oxidase), which ultimately impaired glucose tolerance (Pravenec et al. 2007). Therefore, we decided to compare protein levels in the liver of mtSHR and mtBN strains on both diets by LFQ proteomics. We found that the selective decrease of complex IV is present not only after HFD but also on CHD. Additionally, almost all subunits of complex I were increased on both diets (Figure S4A, B).

The complex IV enzyme activity was reduced in agreement with the suppressed level of complex IV subunits. However, the activity of complex I remained unchanged (Figure S4C). As was previously published, the mtBN strain harbours a unique variant of the *mt-Co1* mitochondrial gene resulting in a single amino acid substitution (Table S1) (Pravenec et al. 2007). Therefore, *mt-Co1* SNP in mtBN strain leads to a decrease of complex IV not only after metabolic challenges (such as a high-fructose or high-fat diet) but also under basal conditions. Further, complex IV level reduction may become rate-limiting under specific circumstances in tissues other than the liver and heart.

### Lower oxidative capacity in the heart of mtF344 rats on HFD is caused by decreased content of OXPHOS complexes

In the next step, we focused on the mtF344 strain and attempted to identify factors responsible for decreased respiratory capacity on HFD. First, we performed LFQ proteomics and analysed proteome-wide responses in the liver and heart, focusing on the OXPHOS apparatus. In the liver, we observed a slight yet significant increase in the overall content of mitochondrial proteins on CHD and no difference on HFD (Figure 6A). Detailed analysis of subunits of OXPHOS complexes revealed a slight rise in some of the OXPHOS subunits in the mtF344 group on CHD but a decrease of complex I and III subunits on HFD (Figure 6C). Decreased abundance of OXPHOS subunits was more pronounced in the heart on HFD, where not only subunits of all OXPHOS complexes were reduced (Figures 6B, C), but even the content of all mitoproteins dropped (Figure 6B). These data indicate that the lower respiratory capacity observed in the heart of mtF344 animals on HFD is caused by reduced OXPHOS proteins.

**Figure 6.**
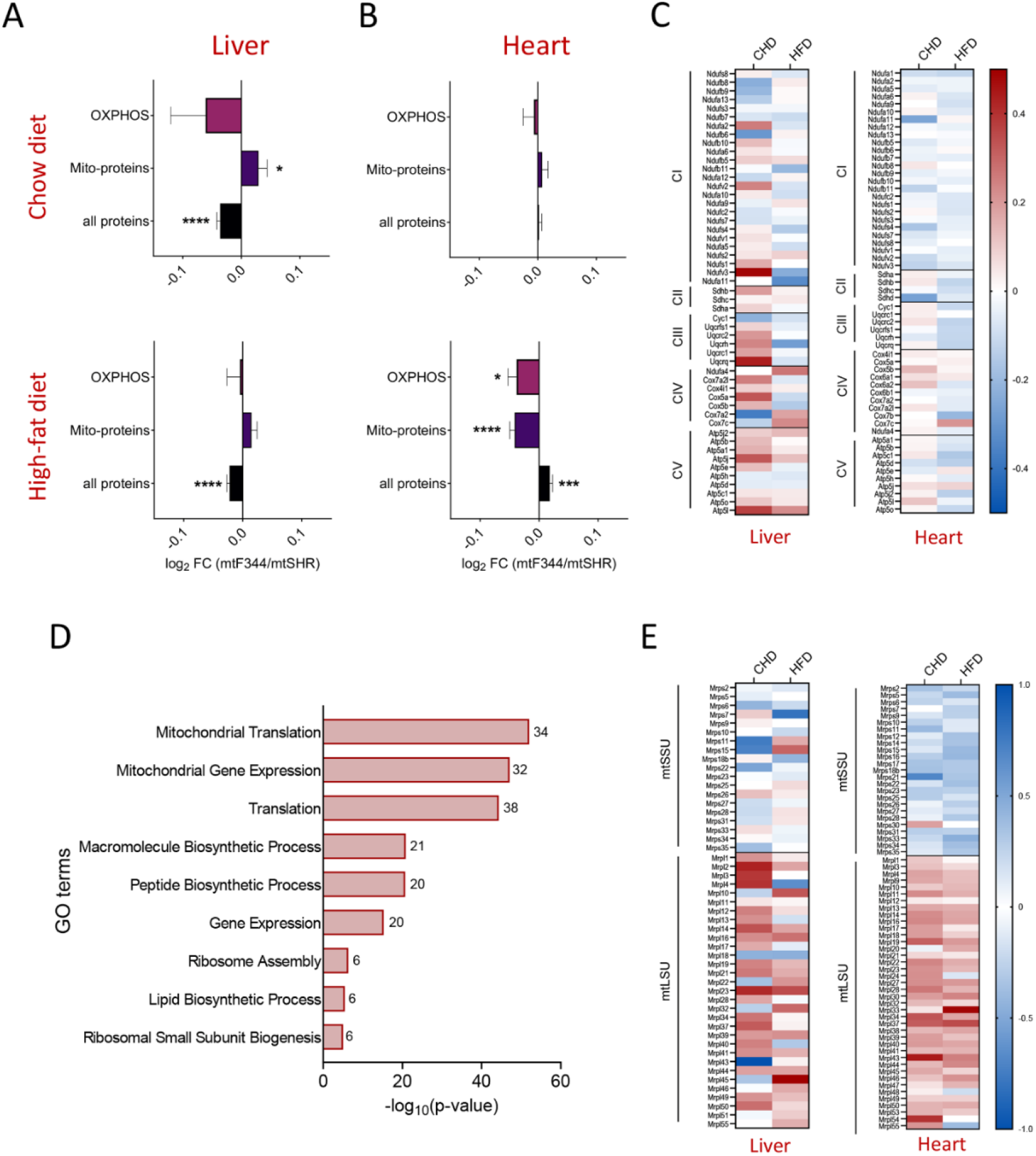
Analysis of protein levels in the liver and heart in mtF344 strain. LFQ-MS analysis of liver (A) and heart (B) of mtF344 compared to mtSHR control group. Data represent mean ± SEM of all proteins, mitochondrial (Mito-proteins – MitoCarta3.0 annotated proteins) and OXPHOS (subunits of complex I–V). (C) Heat maps depicting the average log_2_ fold-change of individual subunits of OXPHOS complexes between mtF344 and mtSHR groups in heart on HFD. (D) GO enrichment analysis using differentially expressed proteins in the heart of mtF344 compared to the mtSHR group on HFD. Numbers represent gene counts in particular GO term. (E) Heat maps depicting the average log_2_ fold-change of individual proteins of mitochondrial small (mtSSU) or large (mtLSU) ribosome subunits. The significances in (A) and (B) were calculated as one sample t-test compared to the mtSHR control group (n = 6). Asterisks represent p-value: * <0.05; ** <0.01; *** <0.001 **** <0.0001.

Supposedly, mitochondrial content is decreased on HFD in the heart of mtF344 animals relative to mtSHR controls. To confirm this hypothesis, we measured mtDNA copy number and citrate synthase activity, established markers of the mitochondrial content in both tissues. As depicted in Figure S5, mtDNA copy number was comparable between genotypes as well as diets in the liver, and citrate synthase was slightly decreased in mtF344 on CHD (Figure S5A). On the other hand, both parameters were suppressed in mtF344 heart tissue (Figure S5B), supporting the results from LFQ analysis.

Subsequently, we checked whether the reduced mitochondrial content resulted from accelerated mitochondrial autophagy. We assessed the autophagosome marker LC3B normalised to citrate synthase in the heart and found that the mitochondrial autophagy was significantly decreased in mtF334 tissue on both diets (Figure S6A).

To explore potential mechanisms behind the OXPHOS decrease in the heart on HFD, we performed a GO enrichment analysis of proteins significantly changed in the LFQ dataset. It identified pathways related to mitochondrial translation as most significantly enriched (Figure 6D). Since the higher levels of N-formylmethionine (fMet) were shown to decrease mitochondrial protein synthesis in a haplotype-dependent fashion in humans (Cai et al. 2021), we assessed fMet levels by untargeted LC-MS approach. We identified significantly higher fMet levels on HFD relative to CHD in mtSHR animals (Figure S6B). However, under the same conditions, the levels of OXPHOS subunits were not significantly decreased (Figures S6C). Furthermore, fMet levels were markedly reduced in mtF344 compared to the mtSHR on HFD (Figure S6B), which again did not correlate with the levels of OXPHOS proteins, which were decreased under the same conditions (Figure 6C). Based on these data, we conclude that fMet levels do not regulate mitochondrial proteostasis in our model system.

Then, we checked the levels of the mitochondrial ribosome proteins by LFQ. We found pronounced decrease in proteins of the small mitochondrial ribosomal subunit (mtSSU) in the mtF344 group, especially on HFD (Figure 6E).

On the other hand, we also observed an increased amount of proteins from the large mitochondrial ribosomal subunit (mtLSU) that was diet-independent in both tissues (Figure 6E). In conclusion, the results implicate that due to the reduced amount of mtSSU, the OXPHOS proteins and mitochondrial content are generally suppressed. Moreover, the cells try to compensate for lower mitochondrial mass by reduced mitophagy, possibly leading to less strict quality control.

### Downregulation of small mitochondrial ribosomal subunit proteins leads to the attenuation of mitochondrial proteosynthesis

Based on these data, we hypothesised that the drop in MRPS quantity may lead to slower synthesis of mtDNA encoded proteins. Thus, we performed *in vivo* metabolic labelling in rat skin fibroblasts derived from mtF344 and mtSHR animals to assess the rate of mitochondrial translation. After 3-hour incubation with DMEM medium containing ^35^S-methionine and ^35^S-cysteine, the amount of newly translated mtDNA encoded proteins was significantly decreased in mtF344 animals (Figure 7A). This decrease affected all evaluated OXPHOS subunits, indicating a general drop in mitochondrial proteosynthesis expected for the system with reduced content of the mtSSU. At the same time, the levels of marker nuclear subunits of individual OXPHOS complexes were not changed (Figure 7B). This suggests that the slower mitochondrial protein translation is still sufficient in skin fibroblasts under non-stress conditions to achieve mitochondrial proteostasis. These results agree with our findings in the animals on a chow diet (i.e. under non-stress conditions), where we did not observe any changes in OXPHOS protein content (Figure 6A–C).

**Figure 7.**
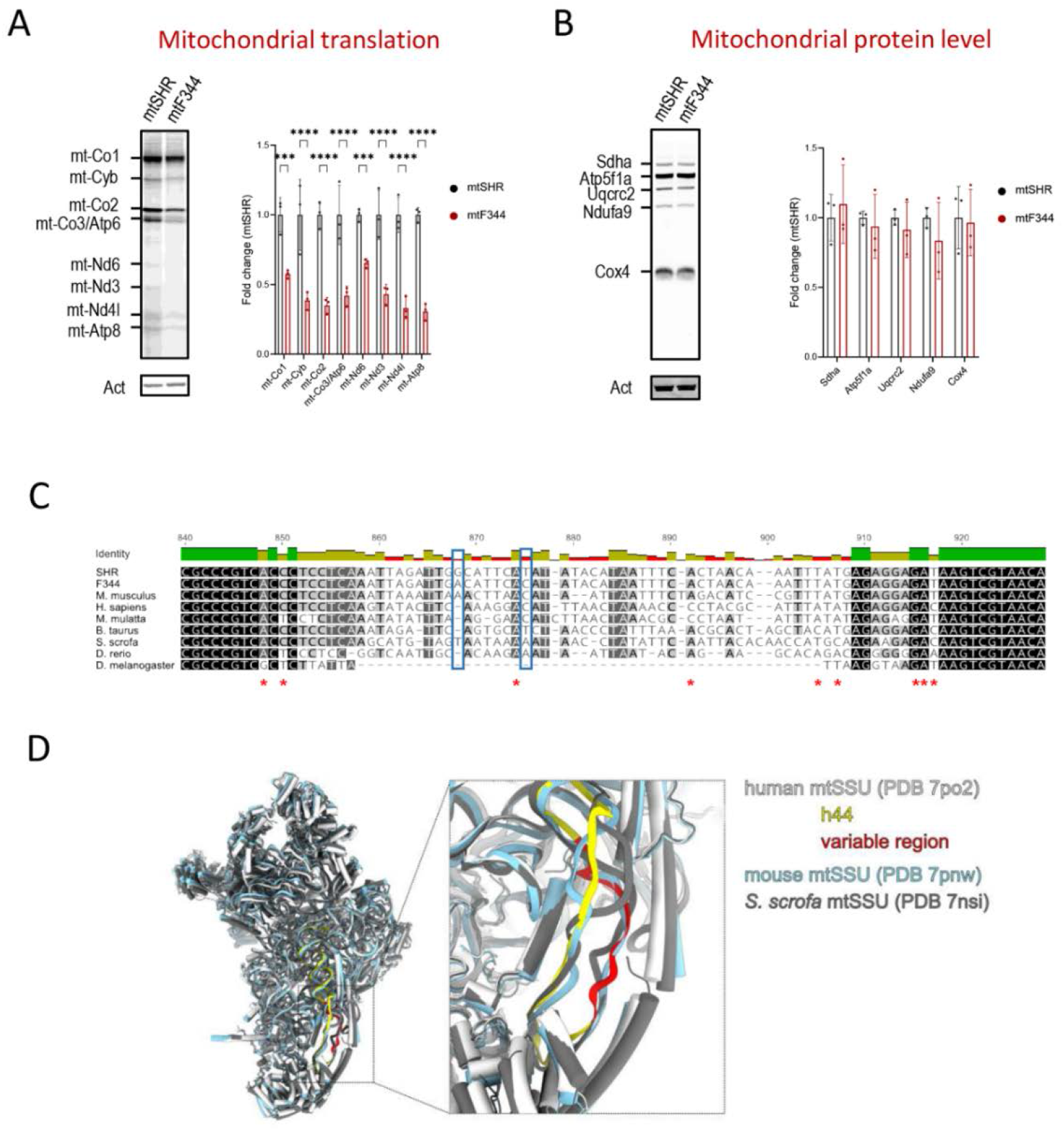
Mitochondrial protein translation and the small mitoribosomal subunit mtSHR and mtF344 strains. Representative images and quantification of three independent samples. (A) Metabolic *in vivo* labelling with ^35^S of mtDNA-encoded OXPHOS subunits after 3h with ^35^S-methionine and ^35^S-cysteine. Representative image shows autoradiographic detection of labelled proteins in 60 μg protein of whole cell lysates analysed by SDS-PAGE. mt-Nd1, 3, 4l and 6 – complex I; mt-Cyb – complex III; mt-Co1–3 – complex IV; mt-Atp6, 8 – complex V. Actin (Act) antibody was used as a loading control. (B) Representative Western blot analysis of the steady-state levels of subunits Ndufa9 (complex I); Sdha (complex II); Uqcrc2 (complex III); Cox4 (complex IV), and Atp5f1a (complex V). Actin IgM (Act) antibody was used as a loading control. Asterisks represent p-value: * <0.05; ** <0.01; *** <0.001 **** <0.0001 (n = 3). (C) Multiple sequence alignment of h44 from selected species was generated by Clustal Omega (Sievers et al. 2011) and visualised in Geneious Prime (Biomatters Ltd.). Sites of single nucleotide polymorphisms in F344 mtDNA are highlighted in blue rectangles. (D) Structures of human (white, PDB 7po2) (Itoh et al. 2022), mouse (pale blue, PDB 7pnw) (Itoh et al. 2022) and porcine (grey, PDB 7nsi) (Kummer et al. 2021) small mitoribosomal subunits (intersubunit side) were superposed and visualised in ChimeraX (Goddard et al. 2018). The helix h44 of human mtSSU rRNA is shown in yellow, and the variable region is red.

Mammalian mtSSU is composed of 30 MRPS proteins and a 12S rRNA, encoded in the mitochondrial genome by the *mt-Rnr1* gene. Interestingly, compared to SHR strain, *mt-Rnr1* of F344 strain possesses two single nucleotide polymorphisms (Table S2). They are localised in the 3’ minor domain helix h44 (Voorhees and Ramakrishnan 2013, Smith et al. 2014), which contains two adenines that are universally conserved in all ribosomes and play a key role in decoding of mRNA. We compared the sequence of helix h44 of *mt-Rnr1* from the SHR and F344 strains and 7 other species (Figure 7C). While the 5’ and the 3’ ends of h44, which form the proximal part of the stem involved in decoding, are highly conserved, the two polymorphisms are in the variable region corresponding to the distal part of the helix. Superposition of human, mouse and porcine mtSSU structures showed that the variable region (depicted in red in Figure 7D) adopts slightly different structures in the three species, and it is relatively loose and exposed to the mtSSU surface on the intersubunit side. During the biogenesis of mtSSU, the variable part of h44 assumes its mature conformation only after dissociation of the assembly factor NOA1, which interferes with its position (Harper et al. 2023). The polymorphisms might affect the process of h44 folding or play a role in an earlier, so far uncharacterized, phase of the assembly (Itoh et al. 2022, Harper et al. 2023). Together, these findings may indicate a possible interaction with yet unknown regulator, either during the translational cycle or during the mtSSU assembly.

We propose that the mtDNA sequence variance in *mt-Rnr1* of the F344 strain may affect mtSSU assembly or a step in the translational cycle, leading to slower mitochondrial protein synthesis and possibly represents a risk factor for insulin resistance and metabolic syndrome development.

In summary, single nucleotide polymorphisms in mitochondrial ribosomal RNA lead to the lower content of the mtSSU subunit of the mitochondrial ribosome. This limits the rate of mitochondrial translation and, under stress conditions (such as a high-fat diet), may cause a drop in the content of OXPHOS complexes. As a result, the overall OXPHOS capacity is compromised, causing inefficient oxidation of substrates including fatty acids, which subsequently leads to the accumulation of bioactive diacylglycerols. Accumulated diacylglycerols may then interfere with insulin signalling and ultimately manifest as an impairment in insulin sensitivity. These disturbances occur in a stress- and tissue-dependent manner, implying the role of protein thresholds across different tissues (Rossignol et al. 1999, Nuskova et al. 2020).

## Discussion

Metabolic syndrome represents a significant burden in developed societies. While its initial trigger in the form of increased body weight may not be a problem, it is associated with other symptoms, including insulin resistance, type 2 diabetes, cardiovascular diseases or cancer (McLaughlin et al. 2017). Notably, common variants of mitochondrial DNA (mtDNA) have been indicated as potential risk factors for symptoms of metabolic syndrome in the human population (summarised in (Ludwig-Slomczynska and Rehm 2022)). However, the mechanism underpinning this link is still largely unknown. Furthermore, there may be numerous pathways linking primary mtDNA sequence variation to metabolic syndrome. Since mtDNA encodes the structural subunits of OXPHOS complexes, primary protein level variation will always stem from them. However, physiological response to variants in complex I may be different to variation at the level of complex III or IV. Furthermore, variability in tRNA or rRNA may affect mitochondrial translation in general and have further reaching consequences.

Due to the variation in nuclear DNA, the impact of naturally occurring single nucleotide polymorphisms (SNPs) in mtDNA in humans may be concealed. To isolate the effect of mtDNA difference, the models of conplastic cell lines (Fang et al. 2018, Chalkia et al. 2018, Cai et al. 2021), conplastic mice (Latorre-Pellicer et al. 2016) and conplastic rats (Pravenec et al. 2007, Houstek et al. 2012, Houstek et al. 2014) possessing identical nuclear genome, but different mtDNA sequence have been developed. Compared to the majority of mouse strains that share the same haplotype (Goios et al. 2007, Pravenec 2013), the inbred rat strains are divided into four major mtDNA haplogroups (Pravenec et al. 2007, Pravenec 2013) – Brown Norway (BN), Fisher F344 (F344), Lewis (LEW) and spontaneously hypertensive rat (SHR) strains. To analyse the contribution of mtDNA variants on complex pathophysiological traits in SHR, conplastic strains possessing the four different mtDNA haplotypes on the SHR nuclear background (mtSHR, mtBN, mtF344 and mtLEW) were derived. The strains differ only in single nucleotide polymorphisms in mitochondrial OXPHOS structural genes, tRNAs and rRNAs with no differences in the nuclear DNA (Pravenec et al. 2007, Houstek et al. 2012, Houstek et al. 2014), summarised in (Pravenec et al. 2021). The mtDNA variation in these strains was associated with a selective reduction in the content and activity of OXPHOS complexes and risk factors for type 2 diabetes (T2D) when the animals were exposed to a high-fructose diet for two weeks (Pravenec et al. 2007, Houstek et al. 2012, Houstek et al. 2014, Pravenec et al. 2021).

Besides sugar-based diets, obesity-inducing diets highly enriched in fats (high-fat diet, HFD) mimic the dietary habits of the human population and therefore are widely used in animal experimental studies (Kleinert et al. 2018). However, the literature focused on the role of naturally occurring mtDNA variations during HFD-induced obesity is limited. Comparison of mice with C57BL/6 nuclear genome and either progenitor mtDNA or mtDNA from NZB/OlaHsd strain revealed the changes in body weight and adipocyte size after three months on HFD (Latorre-Pellicer et al. 2016). Also, distinct mitochondrial genetic background significantly impacted the animal metabolic efficiency in the model of mitochondrial-nuclear exchange (MNX) mouse exposed to HFD for 6 weeks, in which the mtDNA from the C3H/HeN mouse was inserted onto the C57/BL6 nuclear background and vice versa (Dunham-Snary et al. 2018). Further, in the mouse strains that differ in mtDNA encoded subunits of complex I (mt-Nd2 and mt-Nd5), the HFD induced changes in gut microbiota (Kunstner et al. 2022). To provide comprehensive knowledge about the impact of common polymorphisms in mtDNA on metabolic phenotype in obesity, we exposed SHR rats possessing parental (mtSHR), F344 (mtF344) or BN (mtBN) mitochondrial genome to HFD for 15 weeks. In agreement with the data in fructose-fed animals (Pravenec et al. 2007, Houstek et al. 2014), we found decreased glucose tolerance in the mtBN strain and increased insulin level after 30 minutes in mtF344 (Figure 1D, E). Furthermore, increased homeostatic model assessment (HOMA-IR) implies that both conplastic strains are predisposed to impaired insulin sensitivity during obesity (Figure 1F). The mechanisms proposed to explain HFD-induced insulin resistance include increased production of pro-inflammatory cytokines, higher oxidative stress or accumulation of bioactive lipids (Qatanani and Lazar 2007, Sergi et al. 2019). In our model, we identified that mtF344 animals accumulated diacylglycerols (DGs) in plasma, liver and heart (Figure 4B). Increased DGs have been shown to increase phosphorylation of insulin receptor in a protein kinase C dependent manner, which ultimately leads to the development of insulin resistance (Yu et al. 2002). In the mtBN strain, none of the studied mechanisms was altered. However, liver and heart triacylglycerols were increased, indicating dyslipidemia in the tissues (Figure 4C).

The accumulation of lipid intermediates could be caused by inefficient oxidation of metabolic substrates. Indeed, the studies in obese individuals suffering from insulin resistance demonstrated lower oxidative capacity (Kim et al. 2000) compared to healthy controls. It was also shown that overall respiratory chain activity measured as oxidoreductase activity of NADH:O_2_ is reduced in individuals suffering from T2D (Kelley et al. 2002). In the current study, we observed tissue-specific differences in oxidative capacities. In agreement with published data (Raffaella et al. 2008, Cardoso et al. 2013), the fatty acid-dependent (HFD) OXPHOS capacity in the liver of all three groups was significantly increased when palmitoyl carnitine was used as a substrate and attenuated with the mixture of the substrate (compare Figures 5B and C). Since the level of OXPHOS complexes is not changed or decreased after HFD, the data indicate that flux of electrons from fatty acid β oxidation towards the respiratory chain is accelerated. Indeed, the levels of almost all proteins involved in β oxidation in the liver during HFD are elevated (Figures S2).

While the increase of oxidative capacity of fatty acids during HFD is apparent in the liver, the published data addressing this issue in the heart are ambiguous. Using isolated heart mitochondria and palmitoyl carnitine as a substrate, the oxygen consumption was not changed in Wistar rats fed with HFD for three weeks (Cole et al. 2011) but decreased in C57BL6 mice on HFD for 24 weeks (Shao et al. 2020). In the case of our models, diet did not affect neither the level of β oxidation proteins (Figure S3), nor the mitochondrial respiration; however, the oxidative capacity was reduced in mtF344 animals fed with HFD (Figure 5E-G). It indicates deterioration of mitochondrial function caused by lipids overload that leads to DGs accumulation due to inefficient substrate oxidation in the mtF344 strain caused solely by mtDNA sequence variation (summarised in (Sergi et al. 2019)).

Analysis of the respiratory reserve capacity revealed that in the liver, the RS_p_ (the capacity of the respiratory chain that is not utilised during ADP phosphorylation) is comparable to the OXPHOS capacity (Figure 5H). This represents a reserve in the mitochondrial respiratory chain, which may get utilised during a higher substrate supply (e.g. from fatty acids). The RS_p_ was significantly decreased in the mtF344 strain, but apparently it was still sufficient to deal with electrons originating from a higher rate of β oxidation. The RS_p_ was much lower in the heart and did not reflect genotype or dietary intervention (Figure 5I). As the heart uses fatty acids as the primary substrate to produce ATP (Murashige et al. 2020), the data imply that heart mitochondria operate close to the maximal fatty acid-dependent oxidation, thus even a slight reduction of mitochondrial function may translate into observable phenotype.

The different response to HFD in various tissues is also supported by the finding that the biochemical thresholds of OXPHOS complexes differ among the tissues (Rossignol et al. 1999). For instance, the complex IV threshold that reflects excess enzyme activity is the highest in the liver (Rossignol et al. 1999). The detailed analysis of protein levels and activities of OXPHOS enzymes in the mtBN strain demonstrated selective reduction of the complex IV on both diets (Figure S4) and decreased complex IV in mtBN was also observed previously in animals fed a high-fructose diet (Pravenec et al. 2007). Although we did not observe significant changes in the mitochondrial function in the liver and heart of the mtBN strain, the tissue threshold effects may implicate that reduced complex IV activity is limiting in other insulin-sensitive tissues.

In the mtF344 strain, we demonstrated that mitochondrial mass, including OXPHOS proteins, is diminished in the heart of HFD-fed animals (Figures 6B, C and S5B), which was not a consequence of increased autophagy (Figure S6A). GO enrichment analysis of differentially expressed proteins identified mitochondrial translation as the most significantly changed process in the heart of mtF344 fed with HFD (Figure 6D). Recently, changes in circulating N-formylmethionine (fMet) levels were associated with human mtDNA haplogroups. It was demonstrated that higher levels of fMet modulate mitochondrial and cytosolic protein synthesis (Cai et al. 2021). This is of interest, since fMet initiates mitochondrial translation and may be a relevant player in our mtF344 animals. In response to HFD, the levels of fMet were increased in the heart of mtSHR. However, no change in the levels of the subunits of OXPHOS complexes was observed (Figure S6B, C). Moreover, compared to mtSHR, in HFD-fed mtF344 animals, the decreased fMet levels and lower levels of OXPHOS proteins (Figures S6, 6) do not agree with published data (Tucker et al. 2011, Arguello et al. 2018, Cai et al. 2021). Therefore, we concluded that fMet-mediated regulation of protein translation is not the cause of altered mitochondrial translation in mtF344 animals.

Further, the proteomic analysis demonstrated reduced levels of the proteins constituting a small mitochondrial ribosomal subunit (mtSSU) (Figure 6E). Concomitantly, the mitochondrial protein synthesis was attenuated in skin fibroblasts derived from mtF344 animals compared to mtSHR, although the mitochondrial protein level was unchanged (Figure 7A, B). However, these proteins are encoded by nuclear DNA, which is identical in the conplastic strains. In contrast, the rRNA constituent of mtSSU encoded by the *mt-Rnr1* gene harbours two mismatches between mtSHR and mtF344 strains representing the only genetic variability for mtSSU of these strains.

The *mt-Rnr1* polymorphisms identified in mtF344 are located in the variable part of the 3’ minor domain helix h44 (Figure 7C), a key decoding centre component. The maturation of h44 requires dissociation of the NOA1 assembly factor that otherwise hinders the interaction of h44 with the docking site (Harper et al. 2023). Our data showing decreased levels of mtSSU proteins and attenuated mitochondrial protein synthesis are consistent with affected mtSSU biogenesis, likely resulting from the sequence variability in *mt-Rnr1*. Similarly, it was demonstrated that mutations in genes encoding mtSSU proteins usually impair also other mtSSU components, while the proteins of the large subunit of mitochondrial ribosome remain unaltered (Richman et al. 2015, Lake et al. 2017, Borna et al. 2019). Both *MT-RNR1* and *MT-RNR2* represent the most constrained sequences within human mtDNA with highest percentage of invariable bases (Bolze et al. 2020). Therefore, also pathogenic nucleotide substitutions in human *MT-RNR1* are relatively rare, and mostly associate with deafness. Only two (A1555G and C1494T) have been confirmed and several others have reported status in MITOMAP database (Figure 7C, red asterisks) and they are the major contributors to aminoglycoside-induced and non-syndromic genetic hearing loss in patients with maternally inherited DEAFness, autism spectrum intellectual disability, possibly antiatherosclerotic ("MITOMAP: A Human Mitochondrial Genome Database. http://www.mitomap.org"). Transmitochondrial A1555G cell lines pointed to a role for mitochondrial protein synthesis (Guan et al. 2001), but in the absence of experimental models of mtDNA manipulation, the molecular mechanisms involved remained largely unknown (Raimundo et al. 2012, Lee et al. 2015, O’Sullivan et al. 2015). These variants localize to the same h44 helix of *MT-RNR1* as the polymorphisms identified in mtF344 rats, yet predominantly DEAFness polymorphisms involve evolutionary more conserved nucleotides (Figure 7C, D). It is therefore feasible, that mutations at more conserved positions have more severe DEAFness associated phenotype, while those at positions with lower degree of conservation may contribute to metabolic syndrome. Interestingly, there is documented C1518T polymorphism in human population mapping exactly to the same position as C942T polymorphism in mtF344 rats, which occurs at frequency of 10^-4^ to 10^-5^ ("MITOMAP: A Human Mitochondrial Genome Database. http://www.mitomap.org"). However, its potential link to metabolic syndrome would have to be established. Another potential link between *MT-RNR1* sequence variation and diabetic traits was suggested by the mitochondrial genome-wide association mapping of metabolomic phenotypes. By this approach two prominent polymorphisms in *MT-RNR1* were identified (G715A and A856G), which associated with C2/C10:1 or SM (OH)C16:1/lysoPC a C28:1 ratio, respectively (Aboulmaouahib et al. 2022). All these metabolites have links to regulation of insulin secretion and may therefore be analogous to the phenotype we identified in mtF344 rats. Ultimately, it should also be mentioned, that within *MT-RNR1* sequence is encoded 16AA microprotein MOTS-c, which is exported from mitochondria and appears to affect the regulation of cellular metabolism and insulin action in age-related diseases, such as type 2 diabetes mellitus (Kong et al. 2023). However, this mechanism acts independently of mitochondrial protein synthesis.

In summary, the rat conplastic models enabled unique detailed characterisation of physiological variants of the mitochondrial rRNA. The common variation in rRNA sequence leads to a lower rate of mitochondrial translation via the reduction of the mtSSU subunit of the mitochondrial ribosome. Under the stress conditions such as high-fat diet, it results in lower expression of mtDNA encoded proteins and inefficient oxidation of substrates, including lipids. Consequently, accumulated bioactive diacylglycerols may lead to the impairment of insulin signalling. For the first time, we show that the common variants in mitochondrial *mt-Rnr1* may represent a risk factor for insulin resistance.

## Materials and methods

### Animals

Animal experiments were approved by the Institutional Animal Care and Use Committee and the Committee for Animal Protection of the Czech Academy of Sciences (Approval Number: 58/2021) in agreement with the Animal Protection Law of the Czech Republic, which is fully compatible with the guidelines of the European Community Council directives 2010/63/EU. All efforts were made to minimise animal suffering and to reduce the number of animals used. All animals were housed in controlled conditions (22 ± 2 °C, 12:12 hours of light-dark cycle) with access to water and respective diet. The conplastic strains were derived by selective replacement of the mitochondrial genome of highly inbred SHR (spontaneously hypertensive rat) strain with the mitochondrial genome of highly inbred strains of F344 or Brown Norway (53 and 59 backcrosses with SHR males, respectively). The three different strains harbour the SHR (mtSHR), F344 (mtF344) or Brown Norway (mtBN) mitochondrial genome on identical SHR nuclear genetic background (Pravenec et al. 2007, Houstek et al. 2014) (Figure 1A).

### Metabolic phenotype

Each strain was split into two groups of ten individuals, and at the age of 5 weeks (1 week after weaning), we fed one group on a chow diet (CHD, 1310 from Altromin, 14% kcal as fat) and a second group by high-fat diet (HFD, D12492 from Research diets, 60% kcal as fat) for 15 weeks. Body weight and cumulative energy intake (kcal) were estimated once per week. At the end of the study, rats in the *ad libitum* fed state were anaesthetized with isoflurane (2%), blood and tissues were collected for final biochemical analyses, and the animals were sacrificed by cervical dislocation. Blood was centrifuged at 2000 *g* for 10 minutes (room temperature), and plasma was collected and stored at -80 °C.

### Oral glucose tolerance test and insulin measurement

An oral glucose tolerance test (OGTT) was performed at the end of the dietary intervention using a glucose load of 300 mg per 100 g of body weight after 16 h fasting. The glucose was administered by glucose gavage. Blood was drawn from the tail immediately before glucose administration (time point 0) and then after 30, 60, 120, 240 and 360 min timepoints, and the concentrations of glucose and insulin were measured. According to the manufacturer’s instructions, glucose levels were determined using a Glucose (GO) assay kit (Merck). Blood insulin concentrations were measured by Rat Insulin ELISA kit (Mercodia). The homeostasis model for insulin resistance (HOMA-IR) was calculated according to (Antunes et al. 2016) from fasting levels of insulin and glucose by the formula: HOMA IR = serum insulin (pmol/L) *blood glucose (mmol/L)/22.5.

### Gene expression analysis

The total RNA was isolated from the tissues using the RNeasy Plus Universal Mini Kit (Qiagen, Venlo, Netherlands), and cDNA was synthesised from 2 µg of RNA by reverse transcription (SCRIPT cDNA Synthesis Kit, Jena Bioscience GmBH, Jena, Germany). Gene expression assays for *Il1b* (Rn00580432_m1), *Ccl2* (Rn00580555_m1), and *Hprt1* (Rn01527840_m1) were carried out on a ViiA7 instrument (Thermo Fisher Scientific, Waltham, USA) with the following cycling protocols: 95 °C for 12 min and 40 cycles at 95 °C 15 s, 60 °C for 20 s, and 72 °C for 20 s. All reactions were conducted in triplicate, and 1.5 µL of diluted (1:10) cDNA was used in each 5 µL reaction using HOT FIREPol Probe mix Universal (Solis Biodyne, Tartu, Estonia). Transcript quantity was calculated in Quant Studio SW (Thermo Fisher Scientific, Waltham, MA 02451, USA). *Hprt1* transcript levels were used as a housekeeper reference.

### Tissue homogenates and mitochondria isolation

Tissue homogenates (7.5–10%, w/v) were prepared at 4 °C in STE medium (0.25 M (for heart) or in 0.35 M (for liver) sucrose, 10 mM Tris-HCl, 2 mM EDTA, pH 7.4 containing protease inhibitor cocktail, PIC 1:1000, P8340, Merck KGaA, Darmstadt, Germany) using glass-Teflon homogeniser and filtered through a fine mesh (Pecinova et al. 2011). Heart samples were then re-homogenised with Dounce glass-glass homogeniser. All homogenates were used fresh for oxygen consumption measurements or were stored at −80 °C for other assays. For the hydrogen peroxide production assay, freshly liver mitochondria were isolated as described in (Pecinova et al. 2011). Protein content was measured using the Bradford method (Bradford 1976).

### Rat skin fibroblast culture

Primary rat skin fibroblasts were prepared from 3 weeks old mtSHR, mtF344 and mtBN animals, according to (Seluanov et al. 2010). Established cultures of skin fibroblasts were then maintained and subcultured at 37 °C and 5% CO_2_ in air in DMEM medium (Thermo Fisher Scientific) that was supplemented with 10% foetal calf serum (Merck) and penicillin/streptomycin solution (Thermo Fisher Scientific).

### Inflammatory markers

The 10% tissue extracts were prepared from 50 mg of tissue (liver or epidydimal white adipose tissue) by adding 450 μL of extraction buffer (PBS with 0.1% NP-40 and protease inhibitor cocktail 1:500). The extracts were milled in 2 mL tubes containing ceramic bead (30Hz for 3 min) using the mill and centrifuged at 12000 *g* for 10 min at 4 °C. The supernatants were used to assess inflammatory markers Il1b and Ccl2 using LEGENDplex Multiplex assay (BioLegend, San Diego, USA) according to the manufacturer’s instructions.

### Hydrogen Peroxide Production

Hydrogen peroxide production was determined fluorometrically by measuring the oxidation of Amplex UltraRed (Thermo Fisher Scientific) as described in (Pecinova et al. 2017). The assay was performed with 10 μg of liver mitochondria in a KCl-based medium (120 mM KCl, 3 mM HEPES, 5 mM KH_2_PO_4_, 3 mM MgSO_4_, 1 mM EGTA, 3mg/mL BSA, pH 7.2) supplemented with 0.1 mM phenylmethyl sulfonyl fluoride (PMSF) in order to inhibit carboxylesterase that unspecifically converts amplex red to resorufin (Miwa et al. 2016). To distinguish ROS production in different complexes of the electron transport chain, specific substrates and inhibitors were added. Complex I: 10 mM glutamate plus 2 mM malate and 1 μM rotenone, complex II: 0.4 mM succinate or complex III: 10 mM glutamate plus 2 mM malate and 1 μg/mL antimycin A. Amplex UltraRed was used at the final concentration of 50 μM with horseradish peroxidase (HRP) at 1 U/mL. The fluorescence signal from the well containing all substrates and inhibitors, but not mitochondria, was subtracted as a background for every experimental condition used. The signal was calibrated using H_2_O_2_ at the final concentration of 0–5 μM, and H_2_O_2_ stock concentration was routinely checked by measuring its absorption at 240 nm.

### Metabolomics and lipidomics

Metabolomic and lipidomic profiling of plasma, liver and heart samples was performed using a biphasic solvent system comprising of methanol, methyl *tert*-butyl ether, and 10% methanol for metabolite extraction (Sistilli et al. 2021, Hricko et al. 2023). This was followed by four liquid chromatography-mass spectrometry (LC-MS) platforms: (i) analysis of polar metabolites using hydrophilic interaction chromatography in positive ion mode; (ii) analysis of polar metabolites using reversed-phase chromatography (RPLC) in negative ion mode; (iii) analysis of complex lipids using RPLC in positive ion mode and (iv) analysis of complex lipids using RPLC in negative ion mode, with optimised conditions reported previously (Cajka et al. 2023). For data processing, MS-DIAL 4 software was used (Tsugawa et al. 2020), including annotation of polar metabolites using an in-house spectral library combined with MS/MS libraries available from various sources (NIST20, MassBank.us, and MS-DIAL MS/MS library). In addition, complex lipids were annotated using in silico MS/MS spectra available in the MS-DIAL software. Data sets were exported for each matrix and platform as signal intensity from the detector (peak heights) and filtered by removing metabolites with (i) a max sample peak height/blank peak height average <10, (ii) an *R*^2^ <0.8 from a dilution series of quality control (QC) sample, and (iii) a relative standard deviation (RSD) >30% from QC samples injected between 10 actual study samples. The data were then normalised using locally estimated scatterplot smoothing (LOESS) with QC samples injected between 10 actual study samples. The data were analysed in Metaboanalyst 5.0 (Pang et al. 2022).

### Oxygen consumption

The oxygen consumption was measured in liver and heart homogenates as described in (Markovic et al. 2022). Oxygen consumption was measured in the homogenate (0.05–0.15 mg/mL) at 30 °C using the Oxygraph-2k (Oroboros Instruments GmbH, Innsbruck, Austria). The respiratory substrates and inhibitors were used at the following concentrations: 50 μM palmitoyl carnitine, 2 mM malate, 10 mM pyruvate, 10 mM glutamate, 10 mM succinate, 5 μM cytochrome *c*, 1 mM ADP, 5–100 nM oligomycin, 1–3 μM FCCP, 1 μM rotenone, 10 mM malonate. The oxygen consumption rates were analysed using DatLab 5 software (Oroboros Instruments GmbH) and were expressed in pmol O_2_/s/mg protein. Relative spare capacity of ETC to OXPHOS capacity (RS_P_) was calculated as percentage of OXPHOS capacity, i.e. 100*(ETC – OXPHOS capacity)/ OXPHOS capacity.

### Proteomics

Liver samples were pulverised in liquid nitrogen, solubilised in 1% SDS and processed according to the SP4 no glass bead protocol (Johnston et al. 2022). About 500 ng of tryptic peptides were separated on a 50 cm C18 column using a 2.5 h elution gradient and analysed in a DDA mode on Orbitrap Exploris 480 (Thermo Fisher Scientific) mass spectrometer equipped with a FAIMS unit. The resulting raw files were processed in MaxQuant v2.1.4.0. with the label-free quantification (LFQ) algorithm MaxLFQ using the rat proteome database (UP000002494_10116.fasta, UniProt Release 2022_01). Downstream analysis was performed in Perseus (v. 2.0.7.0). GO enrichment analysis of proteins significantly changed in the LFQ dataset was performed by Enrichr software (https://maayanlab.cloud/Enrichr/) (Kuleshov et al. 2016). Data were visualised in GraphPad Prism 9. The mass spectrometry proteomics data have been deposited to the ProteomeXchange Consortium via the PRIDE (Perez-Riverol et al. 2022) partner repository with the dataset identifier PXD045910.

### mtDNA copy number

The total genomic DNA was isolated using the GeneAid Genomic DNA mini kit (Geneaid, New Taipei, Taiwan). The mitochondrial copy number was determined according to (Sadakierska-Chudy et al. 2017), using TaqMan assay for mitochondrial *Nd1* gene and nuclear *Hbb-b2* gene (Rn03296764_s1, Rn04223896_s1, Thermo Fisher Scientific). Quantitative real-time PCR was carried out on a ViiA 7 instrument (Thermo Fisher Scientific, Waltham, MA 02451, USA) using HOT FIREPol Probe mix Universal (Solis Biodyne, Tartu, Estonia).

### Enzyme activities

Complex I and IV activities were determined spectrophotometrically in liver homogenates at 30 °C as cytochrome *c* oxidoreductases (Mracek et al. 2009, Pecinova et al. 2011). The assay medium for complex I contained 50 mM Tris–HCl, 1 mM EDTA, 2.5 mg/mL BSA, 1 mM KCN, pH 8.1 and 100 μM NADH. The reaction was started by adding 40 μM cytochrome *c* and changes of absorbance at 550 nm were monitored. The rotenone-insensitive (5 μM rotenone) portion was subtracted. Complex IV activity was measured in a medium containing 40 mM K-Pi, 1 mg/mL BSA, pH 7.0. The reaction was started with 30 μM reduced cytochrome *c*, and its oxidation was monitored at 550 nm for 40 s. Cytochrome *c* solution (5 mM) was reduced by sodium dithionite. The salt was removed by gel filtration through the Sephadex G-25 column. Citrate synthase (CS) activity was determined in liver and heart homogenates using a medium containing 0.1 M Tris–HCl, 0.1 mM 5,5′-dithiobis-(2-nitrobenzoic acid), 50 μM acetyl coenzyme A, pH 8.1 (Pecinova et al. 2011). The reaction was started by adding 0.5 mM oxaloacetate and then monitoring changes at 412 nm for 1 min. The data were corrected for the absorbance change without oxaloacetate. Enzyme activities were expressed as nmol/min/mg protein using molar absorption coefficient ε_550_ = 19.6 mM/cm (complex I and IV) or ε_412_=13.6 mM/cm (CS).

### Metabolic labelling

The rate of mitochondrial protein synthesis in fibroblast cultures established from individual conplastic rat strains was investigated by metabolic labelling with ^35^S-methionine and ^35^S-cysteine in the presence of emetine, an inhibitor of translation on cytosolic ribosomes, essentially as described (Pravenec et al. 2017, Cunatova et al. 2021). Cells grown to 80% confluency on a 60 mm culture dish were washed three times with PBS. 15 min incubation in DMEM medium without methionine and cysteine was followed by the addition of emetine (100 µg/mL). After 15 min, the medium was exchanged for DMEM medium supplemented with emetine and ^35^S-Protein Labelling Mix (^35^S-Met and ^35^S-Cys, Perkin Elmer NEG072; 100 µCi/mL). Cells were incubated for 3 h at 37 °C, and 250 µM cold methionine and cysteine were added. After 15 min at 37 °C, cells were washed twice with PBS supplemented with 250 µM cold methionine and cysteine, once with PBS and lysed using 150 µL of RIPA solution (150 mM NaCl, 1% Nonidet NP-40, 1% sodium deoxycholate, 0.1% SDS, 50 mM Tris, pH 8.0) supplemented with protease inhibitor cocktail (Merck P8340). Lysates were centrifuged for 15 min at 10000 *g*, and protein concentration in the supernatant was determined by BCA assay (Merck B9643). Finally, 60 µg protein samples were separated using tricine SDS-PAGE (Schagger and von Jagow 1987) on 15% polyacrylamide gels and transferred to a PVDF membrane. The incorporated radioactive methionine and cysteine signal was detected using Typhoon Imager (GE) and quantified using Image Lab 6 software (Bio-Rad). Afterwards, the membrane was probed with a specific antibody against actin (Merck MAB1501) for normalisation of the radioactive signals of mitochondrial proteins.

### Statistical analysis

Data were analysed in GraphPad Prism 5.04 software (GraphPad Software) using unpaired Student’s t-test analysis or one sample t-test (two groups), one-way ANOVA (three and more groups) or two-way ANOVA (for comparisons of more than two parameters). Unless otherwise stated, the data shown are mean values ± S.D. of at least 3 independent experiments. A statistical difference of p < 0.05 was considered significant.

## Acknowledgements

This work was supported by the Czech Science Foundation GACR 19-10354S and the National Institute for Research of Metabolic and Cardiovascular Diseases (Programme EXCELES, ID Project No. LX22NPO5104), and by the grant LUAUS23095 within the INTER-EXCELLENCE program of the Ministry of Education, Youth, and Sports of the Czech Republic. The authors would like to acknowledge the Laboratory of Metabolomics at the Institute of Physiology of the Czech Academy of Sciences and the Proteomics Service Laboratory at the Institute of Physiology (supported by RVO, ID 67985823) and the Institute of Molecular Genetics (supported by RVO, ID 68378050) of the Czech Academy of Sciences.

## Author contributions

Conceptualization, J.H., T.M. and A.P.; Methodology, P.P., M.P, T.M. and A.P.; Formal analysis, M.V., T.Č. and O.G.; Investigation, P.P., K.Č., V.K., G.P.-F., J.Š., T.M. and A.P.; Resources, M.P., T.Č. and M.V., Writing – Original draft, J.H., T.M. and A.P.; Writing – Review & Editing, P.P., K.Č., V.K., K.T., M.V., T.Č., O.G., M.P., J.H., T.M. and A.P.; Visualization, T.M. and A.P.; Funding acquisition, M.P., T.M. and A.P.

## Data availability

All other data described, analysed, and represented in the figures present in this study are available from the corresponding authors upon reasonable request. LFQ-MS data will be made available on PRIDE by the time of publication.

