## Supplementary data for "Physiological variability in mitochondrial rRNA predisposes to metabolic syndrome"

Manuscript title:

Contents:

**Supplementary Tables 1-2**

**Supplementary Figures 1-6**

**Supplementary Figure Legends**

**Supplementary table 1.** Amino acid (AA) substitutions caused by mitochondrial DNA variants in mtSHR, mtF344 and mtBN strains.

| OXPHOS gene | AA No. | mtSHR | mtF344 | mtBN |
|---|---|---|---|---|
| *mt-Nd2* | 18 | Ala | Val | Val |
| | 150 | Ser | Asn | Asn |
| | 244 | Ala | Ala | Thr |
| | 265 | Thr | Ala | Ala |
| | 304 | Met | Thr | Met |
| | 316 | His | 1AA del | His |
| *mt-Nd4* | 23 | Thr | Ile | Ile |
| | 356 | Thr | Ala | Ala |
| | 393 | Ile | Met | Ile |
| | 401 | Ile | Val | Val |
| | 405 | Met | Met | Ile |
| *mt-Nd6* | 30 | Phe | Phe | Leu |
| | 139 | Val | Ile | Ile |
| *mt-Cytb* | 214 | Asn | Asp | Asp |
| | 334 | Val | Val | Ile |
| *mt-Co1* | 2 | Phe | Phe | Leu |
| | 406 | Asn | Asp | Asn |
| *mt-Co2* | 165 | Val | Ile | Val |
| *mt-Atp6* | 35 | Lys | Glu | Glu |
| *mt-Atp8* | 58 | Ile | Ile | Thr |

**Supplementary table 2.** Mitochondrial DNA variants in transfer RNA and ribosomal RNA genes in mtF344 and mtBN strains compare to mtSHR. Nucleotide No.: position in mtF344/position in mtBN (position in mtSHR). NC – no change in sequence with respect to mtSHR sequence.

| | gene | Nucleotide No. | mtF344 | mtBN |
|---|---|---|---|---|
| mt-tRNA | *Phe* | 32 | ins A | ins A |
| | | 52 | del A | del A |
| | *Cys* | 5196/5200 (5198) | A>G | A>G |
| | | 5198/5202 (5200) | A>G | A>G |
| | | 5233/5237 (5235) | T>A | T>A |
| | *Tyr* | 5265/5269 (5267) | G>C | G>C |
| | *Asp* | 6974/6978 (6976) | G>A | G>A |
| | *His* | 11542 (11540) | NC | G>A |
| | *Thr* | 15329 (15331) | G>A | NC |
| | *Pro* | 15400 (15398) | NC | T>C |
| mt-rRNA | *mt-Rnr1* | 648 | NC | A>C |
| | | 935 | A>G | NC |
| | | 942 | C>T | NC |
| | *mt-Rnr2* | 1099 | NC | T>C |
| | | 1130 | ins C | NC |
| | | 1136/1137 (1137) | A>C | A>C |
| | | 1222 (1223) | A>G | NC |
| | | 1247 (1248) | C>T | NC |
| | | 1520/1521 (1521) | G>A | G>A |
| | | 1584/1585 (1585) | T>C | T>C |
| | | 1653-4 | del AC | NC |
| | | 1693 | NC | C>T |
| | | 1717/1716 (1716) | T>C | T>C |
| | | 1833 (1832) | G>A | NC |
| | | 1918 | NC | G>A |
| | | 2171 (2170) | T>C | NC |
| | | 2647 | NC | del T |

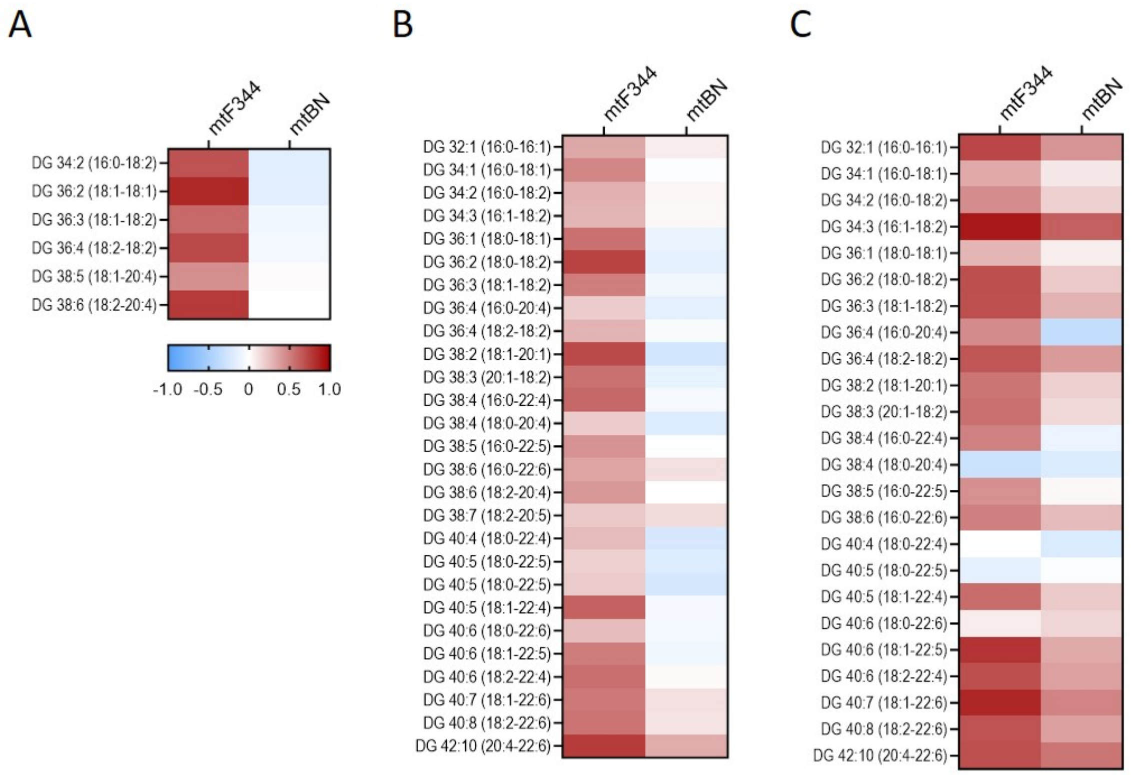


**Supplementary figure 1: LC-MS analysis of individual diacylglycerols in the conplastic strains on HFD.** The heatmap represents average $\log_2$ fold change of individual diacylglycerols compared to mtSHR control in plasma (**A**), liver (**B**), and heart (**C**) of mtF344 and mtBN strains (n = 6).

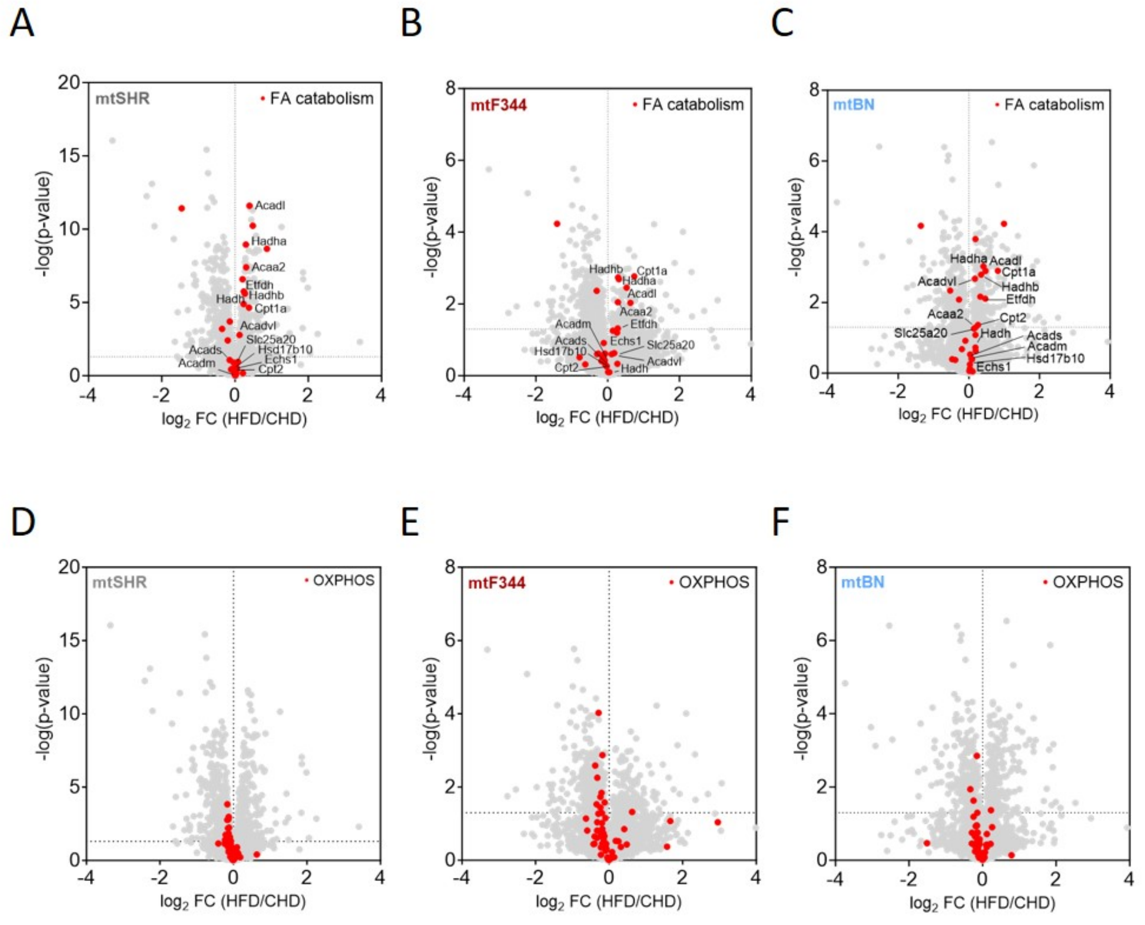


**Supplementary figure 2: Label-free quantification mass spectrometry analysis (LFQ-MS) of liver tissue**. Volcano plots depict differential contents of all proteins in between HFD and CHD in mtSHR (**A, D**), mtF344 (**B, E**) and mtBN (**C, F**). Red circles represent proteins involved in fatty acid (FA) degradation (**A-C**) and OXPHOS (**D-F**) according to KEGG pathways, enzymes of mitochondrial β-oxidation are labeled. Each point represents the data from 6 animals.

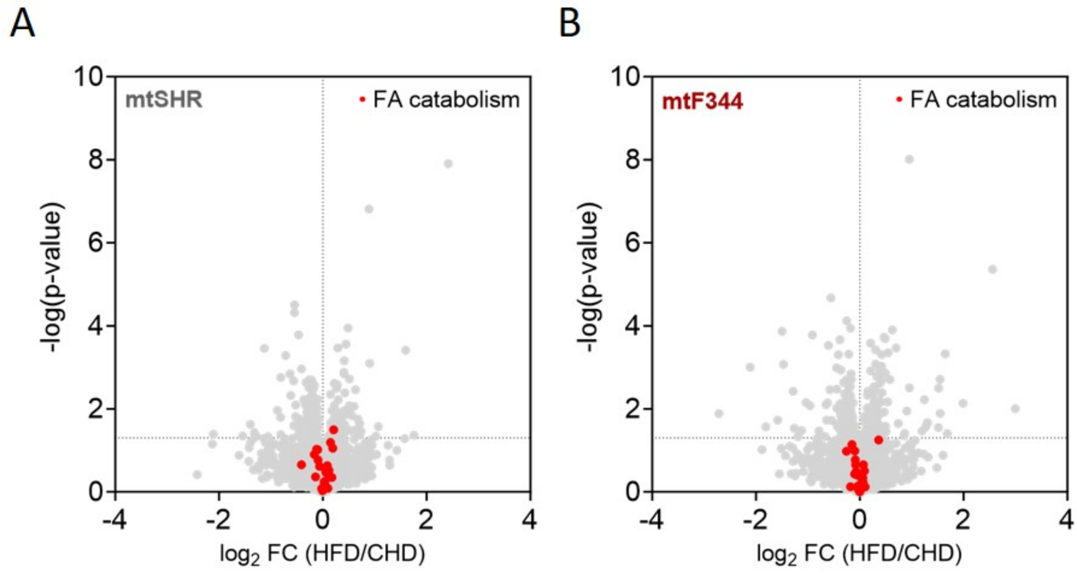


**Supplementary figure 3: Label-free quantification mass spectrometry analysis (LFQ-MS) of heart tissue.** Volcano plots depict differential contents of all proteins in between HFD and CHD in mtSHR (**A**) and mtF344 (**B**) strains. Red circles represent proteins involved in fatty acid (FA) degradation (according to KEGG pathways), enzymes of mitochondrial β-oxidation are labeled. Each point represents the data from 6 animals.

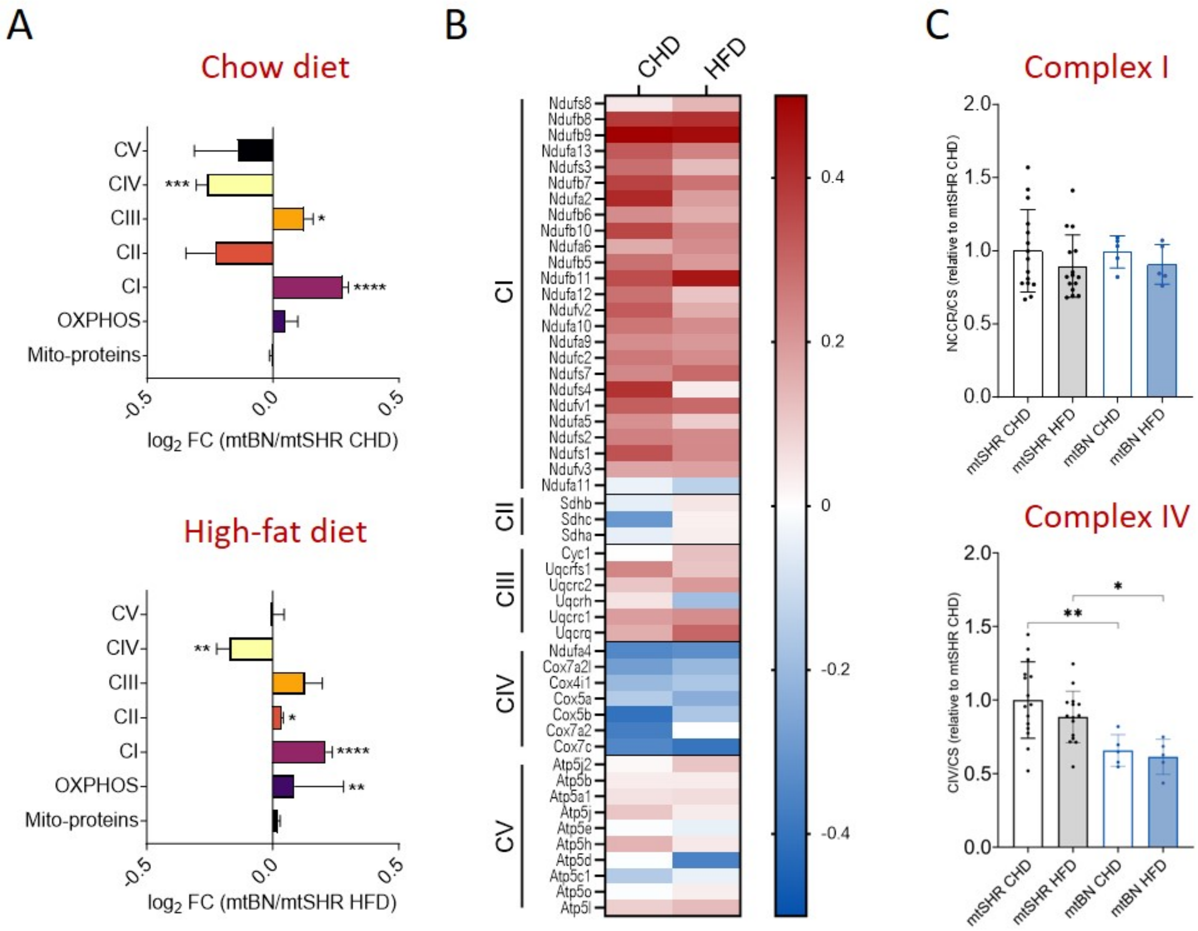


**Supplementary figure 4: Analysis of protein levels in mtBN strain liver.** (**A, B**) LFQ-MS analysis of liver tissue of mtBN compared to mtSHR control group (n = 6). (**A**) Bar graphs depict the average differential content ($log_2$ difference) ± SEM between the strains on chow or high-fat diet. Mito-proteins - mitochondrial proteins annotated in MitoCarta3.0; OXPHOS – proteins of the oxidative phosphorylation; CI-CV – OXPHOS complexes I–V. (**B**) Heatmap of differential content ($log_2$ difference) of individual subunits of OXPHOS complexes. (**C**) The activity of complex I and IV in liver homogenates measured by spectrophotometry and normalized to activity of citrate synthase (CS). Bar graphs depict average enzyme activity ± SD (n = 5). The significances in (**A**) were calculated as one sample t-test compared to the mtSHR control group. Bar graphs represent means ± S.D. from at least 5 animals. Asterisks represent p-value: * <0.05; ** <0.01; *** <0.001 **** <0.0001.

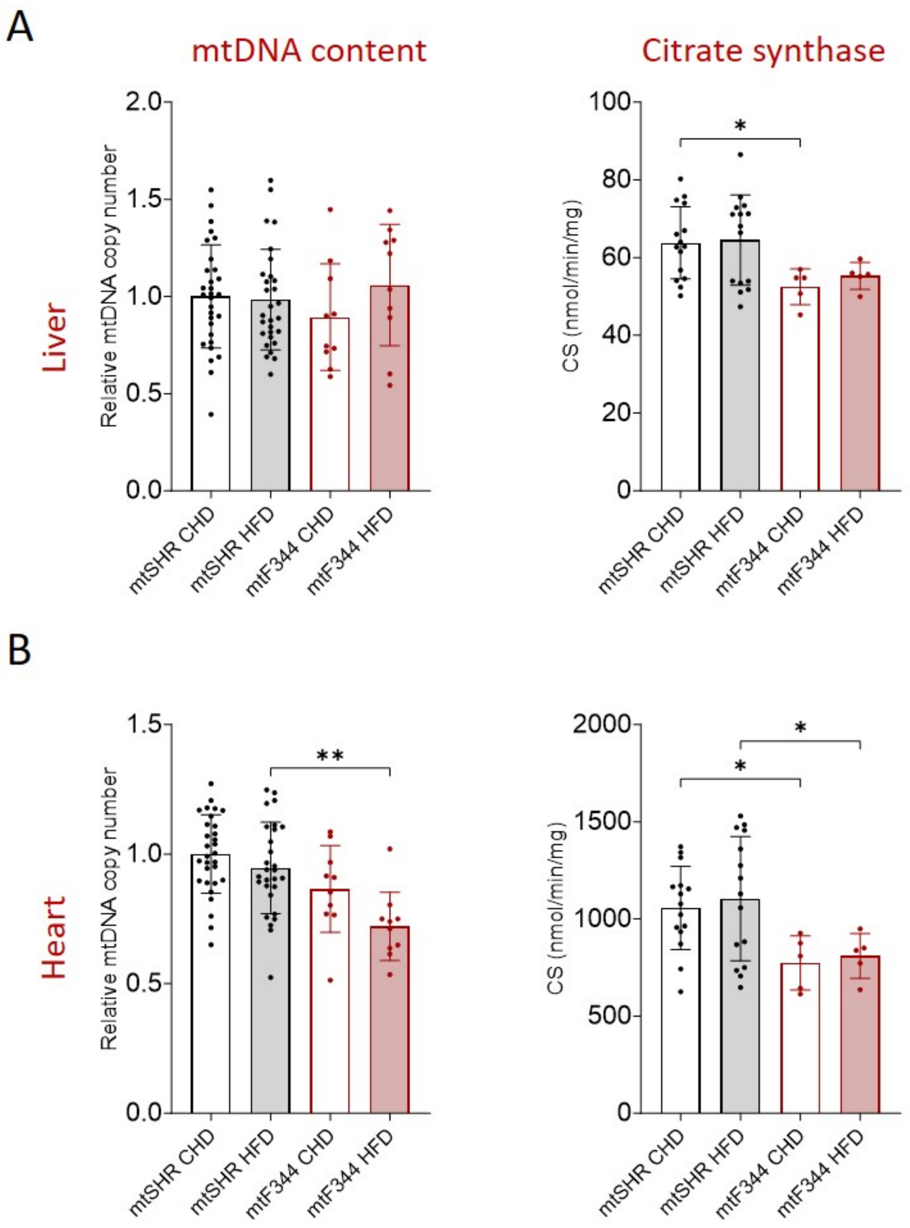


**Supplementary figure 5: Content of mitochondria in mtSHR and mtF344 animals.** Mitochondrial DNA copy number was measured by qPCR and citrate synthase activity (TCA cycle enzyme) was measured spectrophotometrically in liver (**A**) and heart (**B**) tissues. The data are expressed as means ± S.D. from at least 10 animals.

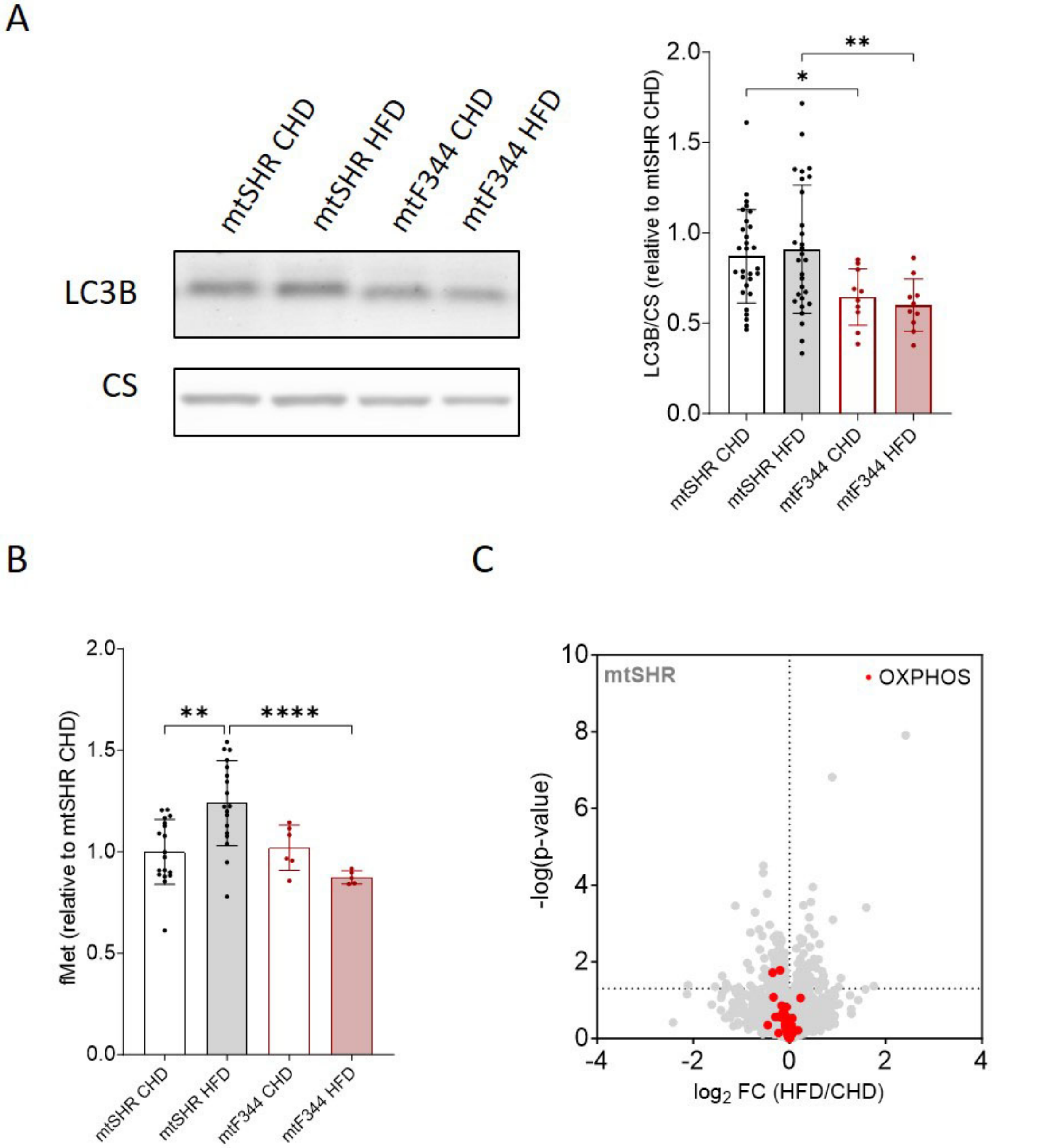


**Supplementary figure 6: Analysis of autophagy and mitochondrial translation initiation in heart tissue of mtSHR and mtF344 animals.** (**A**) Representative Western blot and quantification of LC3B protein in heart homogenates fo at least 10 animals. Graph represents quantification of LC3B normalized to citrate synthase (CS) that was used as a marker of mitochondrial content. (**B**) The level of N-formylmethionine (fMet) in heart tissue measured by LC-MS metabolomics. The data are expressed as means ± S.D. from at least 6 animals. (**C**) Label-free quantification mass spectrometry analysis (LFQ-MS) of mtSHR heart tissue. Volcano plots depict differential contents of all proteins in between HFD and CHD. Red circles represent subunits of OXPHOS complexes. The data are expressed as means ± S.D. and each point represents the data from 6 animals.
